# Genome-wide association identifies impacts of chlorophyll levels on reproductive maturity and architecture in maize

**DOI:** 10.1101/2022.11.07.515492

**Authors:** Rajdeep S. Khangura, Gurmukh S. Johal, Brian P. Dilkes

## Abstract

The semi-dominant mutant allele *Oy1-N1989* encodes a dominant-negative subunit I of Mg-Chelatase that catalyzes the first committed step of chlorophyll biosynthesis. Analyses of mutant F1 hybrids from crosses of *Oy1-N1989* to diverse maize lines identified a natural variant at *oil yellow1* as major modifier of chlorophyll content. Chlorophyll reduction increased the time to reproductive maturity, decreased stalk width, and had complex effects on plant height. We explored these effects by genome wide association studies of F1 hybrids crosses of *Oy1-N1989* to a maize diversity panel. The delay in reproductive maturity in *Oy1-N1989*/+ mutants neither altered the vegetative-to-adult phase change nor total leaf number but did slow the rate of organ emergence from the whorl. In addition to the allele at *oy1*, analysis of tetrapyrrole, heme, and chlorophyll biosynthesis pathway genes identified multiple modifiers of mutant traits. Consistent with organ emergence, rather than a developmental delay, affecting the reproductive delay of *Oy1-N1989*/+ mutants, known flowering time regulators did not affect variation in the reproductive delay affected by chlorophyll loss. Plant height showed a complex relationship with chlorophyll. The *Oy1-N1989*/+ mutant F1 plants with a modest reduction in chlorophyll were taller than their wildtype siblings, but F1 plants with dramatically decreased chlorophyll were shorter than their wildtype siblings. Perturbations in the biosynthesis of the phytochrome chromophore increased plant height in wild-type plants but dramatically reduced plant height and chlorophyll contents of *Oy1-N1989*/+ mutants. The mechanism by which the interaction between chlorophyll and bilin affects height and chlorophyll remains to be explored.

**One sentence summary:** Chlorophyll reduction in maize delays reproductive maturity by altering growth and not development, and depending on the reduction can either increase or decrease plant height.

## Introduction

Photosynthesis is a biochemical process that converts light energy into chemical energy. This process occurs in two sequential steps. The first step is light-dependent, referred to as the light reactions, which pump protons across a membrane-stabilized gradient, evolve O_2_, and ultimately generate ATP and reducing power via NADPH. The second set of reactions is light-independent, referred to as the dark reactions, resulting in a CO_2_ reduction (Calvin and Benson 1948), which ultimately yields glucose through a series of enzymatic steps. This stored energy is primarily transported as sugar-captured carbon from the green parts of the plant to various tissues for metabolism and as the building blocks for growth and biomass. The growth of plant organs is coordinated to use these resources optimally and ensure plant fitness. In maize, interfering with the transport of the energy stored as sugar-captured carbon not only alters plant growth but also delays reproductive maturity (Braun *et al*. 2006; Slewinski *et al*. 2009; Khangura *et al*. 2020a; b)

Photosynthesis can be altered by interfering with enzymes in the core biochemical pathway or the levels of chlorophyll (Long *et al*. 2006, 2022; Khangura *et al*. 2019, 2020a). As chlorophyll becomes limited, varying the level of chlorophyll results in corresponding changes in sugar and starch levels that impacts plant growth, reproductive maturity, and development (Khangura *et al*. 2020a; b). Chlorophylls are porphyrin derivatives required at the reaction center for the conversion of light energy into chemical potential. In porphyrin metabolism, protoporphyrin IX is at the branch between the heme and chlorophyll pathways. The first committed step to chlorophyll is the chelation of a Mg^2+^ ion, catalyzed by the trimeric enzyme Magnesium Chelatase (MgChl). Subunit I of MgChl is encoded by the *oil yellow1* (*oy1*) gene in maize (Sawers *et al*. 2006). A semi-dominant allele of *oy1* called *Oy1-N1989* is encoded by an L161F change that results in pale-yellow seedlings in heterozygous condition (Sawers *et al*. 2006). The same missense mutation in this subunit in other plants and blue-green algae also results in a semi-dominant mutant allele (Hansson *et al*. 1999, 2002). Maize growth directly depends on photosynthetic output, and reduction in the levels of chlorophyll affected by *Oy-N1989*/+ delays reproduction, reduces stem thickness, and alters the height of maize plants (Khangura *et al*. 2020a; b).

We previously identified natural variation in diverse maize lines that modifies the expression of *Oy1-N1989*/+ mutant phenotypes (Khangura *et al*. 2019, 2020a; b). The strongest modifier, *very oil yellow1* (*vey1*), was associated with an expression level polymorphism at *oy1* itself. This affected cryptic variation in chlorophyll accumulation that only impacted phenotypes in the presence of the *Oy1-N1989* allele (Khangura *et al*. 2019). Alleles of *vey1* associated with less wildtype OY1 transcript accumulation exhibited reduced chlorophyll levels in *Oy1-N1989*/+ mutant plants. Reduced chlorophyll levels altered plant morphology and delayed reproductive maturity (Khangura *et al*. 2020a; b). Stalk width was reduced in mutants with lower chlorophyll levels, but the relationship between chlorophyll and plant height was complex (Khangura *et al*. 2020b). The chlorotic *Oy1-N1989*/+ mutants were taller than their wildtype siblings when partially suppressed by the *vey1^B73^* allele, but the mutants were shorter than their wildtype siblings when enhanced by the *vey1^Mo17^* allele. This identified a special case of epistatic interaction between *Oy1-N1989* and the modifier alleles. The alleles at *vey1* did not affect the wild-type plants. *Oy1-N1989* is epistatic to *vey1*, yet the direction of the effect of *Oy1-N1989* on height was dependent on the allele at *vey1. Oy1-N1989*/+ plants were taller or shorter than their wild-type siblings, depending on the *vey1* allele they encoded (Khangura et al., 2020b). There is no known mechanism by which a reduction in chlorophyll increases growth or elongation, and certainly no explanation for a decrease in chlorophyll, making plants taller and shorter. A discontinuity in the accumulation of chlorophyll in the mutant progenies segregating in RIL x *Oy1-N1989*/+ F1 population with the divergent *vey1* alleles prevented distinguishing between the two models. One possibility is that the alleles at *vey1* reversed the effect of reduced chlorophyll on plant height. The second is that a modest reduction in chlorophyll increases plant height by an unknown mechanism, but drastic reductions in chlorophyll cannot support sufficient anabolic metabolism for plant growth. We favored a “goldilocks” model where an optimal chlorophyll accumulation exists at which plant height can be maximized by a heretofore unknown connection between plant height and reduced chlorophyll contents (Khangura *et al*. 2020b).

Quantitative genetic studies can transcend the boundaries of their study system if they link natural variation to molecular causality. We have attempted to do this in our studies of *Oy1-N1989* modifiers. One of the difficulties in assigning molecular causality to alleles identified by studies of natural variation is the divergent concepts and language used to describe the effects of genes and alleles in the quantitative and molecular genetic senses. By comparing the effects of alleles in a diverse population of maize in combination with *Oy1-N1989*, we detected variation affecting chlorophyll levels and could identify the molecular mechanism at this locus. Whole genome sequences provided the molecular nature of all polymorphisms at *oy1* allowed us to compare, at single-nucleotide resolution, the alleles segregating in the maize association panel for roles in *Oy1-N1989* suppression. Lastly, public collections of expression data in both the B73 x Mo17 RILs (Li *et al*. 2013, 2018) and a maize association panel (Kremling *et al*. 2018) allowed eQTL to be assessed as possible mechanisms. Thus, the alleles could be compared with explicit and direct connections to their molecular identities, and the suitability of hypotheses of protein-coding changes versus expression variation could be compared. Having done this with *vey1*, we realized that combining expression data, allelic state, formal hypotheses for mechanism, and allele effect direction could be used to efficiently eliminate candidates from consideration for QTL more generally (Khangura *et al*. 2020b).

In the current study, we used a population of three hundred and forty-three F1 families derived from crosses between a maize association panel and an *Oy1-N1989*/+ tester in the B73 background. Association mapping using these F1 families identified *vey1* as the major modifier of reproductive maturity and stalk width in the *Oy1-N1989*/+ mutants. We extended our approaches to exploring the molecular mechanism of natural variation by conducting pathway-level analyses of enzymes in the tetrapyrrole and chlorophyll biosynthetic pathway. Multiple enzymes in these pathways encoded alleles that modified traits in the *Oy1-N1989*/+ mutants. A similar analysis using known flowering time regulators of maize identified no role for these genes in the *Oy1-N1989*-dependent reproductive delay. Instead, the delay in reproductive maturity in the *Oy1-N1989*/+ mutant results from slower growth affecting organ emergence and not a change in development. Using a continuous range of chlorophyll content in the F1 *Oy1-N1989*/+ hybrids in the association panel, we demonstrate our previous observation of both stimulatory and inhibitory effects of the *Oy1-N1989* allele on plant height due to the existence of optimal chlorophyll levels that underlie the seesaw-effect on plant height. We further used assessments of alleles, rather than loci, to compare mapping results between RIL and GWA experiments. We identified candidate genes within overlapping loci from different mapping studies and populations using allelic state and its effect direction as an additional layer of shared causation between QTLs. Reliance on allelic effects for candidate gene discovery can be used with currently available tools for QTL mapping and GWA. A broad acceptance of this approach would improve inference sharing across experiments and help identify molecular identities underlying natural allelic variations.

## Materials and Methods

### Plant materials

The maize mutant allele *Oy1-N1989* was obtained from the maize genetics COOP (University of Illinois, Urbana/Champaign) in a mixed genetic background and introgressed into the B73 inbred background, as described previously (Khangura *et al*. 2019). The mutants used here were backcrossed eight times to B73 (*Oy1-N1989*/+:B73) using mutant heterozygotes as pollen-parents. For association mapping, a collection of 343 diverse maize inbred lines, described previously (Khangura *et al*. 2019), were selected as a mapping population. These lines were used to make F1 hybrid progenies by crossing them as ear parents to *Oy1-N1989*/+:B73 as a pollen parent, except for dent-sterile popcorn lines where the mutant was used as an ear-parent (**Figure 1**).

**Figure 1.**
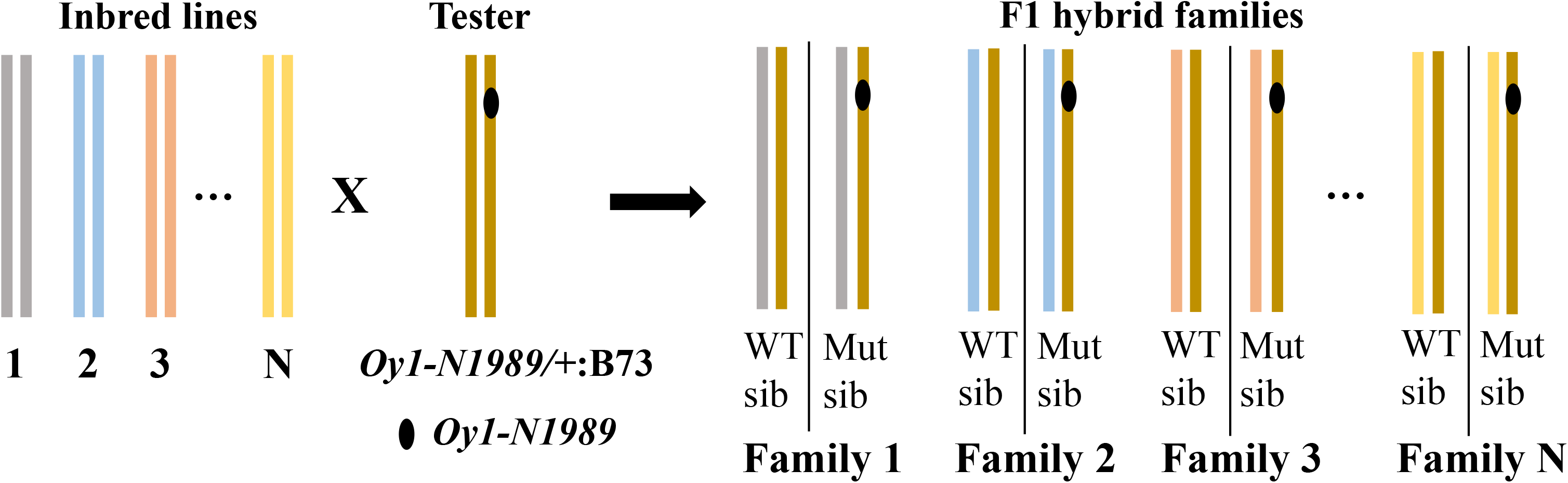
Schematic of the crossing scheme used to create the F1 progenies between the maize inbred lines of the association panel and the heterozygous *Oy1-N1989*/+ tester in the B73 genetic background. The two vertical lines for each inbred, tester, and the F1 hybrid family represent genotypic state for chromosome 10 (homozygous for inbred and tester, and heterozygous for F1 hybrids). The *Oy1-N1989* mutant allele is shown by a black oval. Each F1 hybrid family consists of isogenic wild-type (WT sib) and a *Oy1-N1989*/+ mutant (Mut sib) sibling.

B73-Mo17 NIL lines (Eichten *et al*. 2011) were previously selected so that a set of B73-like and Mo17-like NILs with the alternative allele at the *vey1* locus (Khangura *et al*. 2019). NIL were crossed to *Oy1-N1989*/+:B73 tester to make F1 hybrids to perform a single locus test of *vey1*. Seeds of *elm1-ref* homozygotes were obtained from the maize genetics COOP in the B73 inbred background (stock# 804F). Populations segregating for single and multi-locus combinations of *elm1-ref* and *Oy1-N1989* were generated in the B73 background. To generate this material, pollen from *Oy1-N1989*/+ mutants in the B73 background was used to cross-pollinate ears of *elm1-ref* plants. The progeny of this cross segregated for *Oy1-N1989/+;elm1-ref*/+ and *+/+;elm1-ref*/+ plants in the next generation. Ears of *Oy1-N1989/+;elm1-ref*/+ F1 individuals were crossed to *elm1-ref* homozygotes. The progeny of this second cross segregated 1:1:1:1 for four categories of leaf color: normal/wildtype (+/+;*elm1-ref*/+), pale-green thin (+/+;*elm1-ref/elm1-ref*), yellow-green (*Oy1-N1989/+;elm1-ref*/+), and severe yellow-green (*Oy1-N1989/+;elm1-ref/elm1-ref*).

### Experimental Design

Experiments were conducted at the Purdue Agronomy Center for Research and Education in West Lafayette, Indiana. The test cross progenies of the maize association panel and *Oy1-N1989*/+:B73 were planted in the summer of 2016. The test cross progenies of the NIL and *Oy1-N1989*/+:B73 were planted in the summer of 2019. All F1 families were planted as a single plot of 18 seeds (plant density of ~24,017 plants per acre). Three replications of each F1 family were planted in a randomized complete block design. The F1 progeny of B73 and Mo17 crossed to *Oy1-N1989*/+:B73 were used as randomized checks in each range treated as a block. Height, stalk width, and days to reproductive maturity were measured across all three replications. The average plot length of all experiments described here is 3.84 m with an interrow spacing of 0.79 m and an alley space of 0.79 m. The *elm1-ref* and *Oy1-N1989* mutant population were planted as contiguous field rows without any alley in the summer of 2020. No irrigation was necessary as rainfalls were uniformly distributed around the growing season. Nutrient, pest, and weed management for growing field maize in Indiana were followed.

### Phenotypic measurements

All the F1 progenies resulting from crosses between inbred lines or NILs with *Oy1-N1989*/+:B73 mutants segregated ~1:1 for mutant and wildtype F1 hybrid isogenic siblings. Phenotypes were measured on both mutant and wild-type siblings. Height and stalk width traits were measured ~3 weeks after flowering from roughly three random plants/genotypes within a given plot. For height traits, the vertical distance from the soil line to the point of attachment of the flag-leaf and primary ear node was used to determine the flag-leaf height and ear height, respectively. Ear-to-flag-leaf height was determined by subtracting ear height from flag-leaf height. All primary height measurements are expressed in centimeters (cm). The stalk width of mature plants was measured using Vernier caliper (Vernier 1631) along the shortest diameter of the internode above the primary ear. These measurements were recorded as caliper units, with one caliper unit ~1/32 of an inch or ~0.08 cm. The date of ~50% anthesis and ~50% silking of each genotype within the plot was recorded by walking the field plots every other day during flowering. The date of sowing was subtracted from the date of 50% anthesis and the date of 50% silking to obtain days to anthesis and days to silking, respectively. In addition to using primary phenotypes for mapping, we also calculated differences between wildtype and mutant trait values to obtain difference traits and divided mutant to wildtype values to calculate ratiometric traits. Chlorophyll contents were measured in the *elm1-ref* and *Oy1-N1989* mutant populations using a CCM-200 plus (Opti-Sciences, Inc., Hudson, NH), and the data were expressed as chlorophyll content index measurements (CCM). CCM was measured for the leaf lamina of the top fully expanded leaf at 36 days after sowing. Plant height, stalk width, and days to reproductive maturity were measured in this material as described above. To calculate the rate of vegetative development in BM-NILs x *Oy1-N1989*/+:B73 F1 progenies, the days to a given developmental stage were recorded by periodically visiting the plots for both mutant and wild-type siblings.

### Statistical phenotypic analyses

The preliminary data analysis and visual inspection of the data were done using JMP 15, and outliers were identified and set to missing. The average trait value for each phenotype was obtained within each plot. Analysis of variance determined replication as a significant source of variation across most phenotypes. To account for sources of variation, we used a mixed linear model implemented in R package *lme4* to calculate BLUP. The *lmer* function was used to fit the model where pedigree, replication, and interaction terms were used as random effects. The broad-sense heritability on the line mean basis was calculated from these variance estimates as described by Holland *et al*. 2003. Pairwise differences between genotypes were analyzed by Student’s t-test (Student 1908).

### Genome-wide association

We used 343 inbred maize lines to make F1 hybrids with the *Oy1-N1989*/+:B73 mutant tester. The marker genotypes for 304 inbred lines used for association mapping were obtained from the HapMap3 project (Bukowski *et al*. 2018). Lower-density SNP genotypes were obtained for 39 accessions from a previously published Ames panel dataset (Romay *et al*. 2013). We imputed SNP genotypes for these 39 inbred lines using the HapMap3 positions for the larger panel using the Beagle version 5.1 software package (Browning *et al*. 2018). Imputations were performed using a window size of 12 Mb with a window overlap of 2 Mb, 10 burning, 12 iterations, and an effective population size of 1000.The fully imputed SNPs across all 343 inbred lines comprised 55,242,281 SNP positions. These were filtered to retain positions with a minor allele frequency greater than 0.05. This filtering retained 22,507,869 SNPs that were used for GWA. A random subset of 2% of the SNP was used to calculate principal components and a kinship matrix. We used five principal components and the kinship matrix to control for the population structure and familial relatedness, respectively, in our GWA analyses. GWA analyses were carried out using the compressed mixed linear model (CMLM) implemented in GAPIT package (Lipka *et al*. 2012). Post GWA, the multiple-test correction was done using a Bonferroni-corrected threshold at α=0.05 corresponding to a P-value of 2.2 × 10^−9^ to identify statistically significant associations. After determining the multiple test-corrected associations, we sought to analyze SNP associations with genes of known function at lower statistical significance to avoid false negative conclusions due to a conservative genome-wide threshold. For further analyses, we retained SNP associations at an arbitrary threshold of P-value = 10−4. These associations were used for various pathway and developmental level candidate gene association analyses described in the results section.

### Data availability

The BLUP values of all the traits collected from the F1 hybrids between the maize diversity panel and the *Oy1-N1989*/+:B73 mutant are provided in **Supplemental Table S1**. The dummy genotypes with reference allele coded as “A” and alternate allele coded as “T” is also provided as **Supplemental File S1**.

## Results

### GWA mapping detected *vey1* as a major modifier of reproductive maturity in *Oy1-N1989* mutants

Previously, we detected a strong modifier, *very oil yellow1* (*vey1*), that impacts chlorophyll levels in the *Oy1-N1989*/+ mutants. The *vey1* locus is encoded by a cis-eQTL at the *oy1* gene itself (Khangura *et al*. 2019, 2020a; b). The *vey1* alleles associated with reduced expression of the wildtype allele at *oy1* resulted in a reduction of chlorophyll in *Oy1-N1989*/+ mutants. This Locus was detected by crossing *Oy1-N1989*/+ mutants to RIL populations, NIL, and a maize diversity panel for GWAS (Khangura *et al*. 2019, 2020a; b). In these experiments, F1 families segregating 1:1 for the *Oy1-N1989* and wildtype *oy1* allele are generated, allowing comparison of mutant and wildtype controls within each plot. We have previously demonstrated that chlorophyll content variation in *Oy1-N1989* F1 crosses with B73 x Mo17 RIL modify a suite of other traits, including time to reproductive maturity, plant height, and stalk width. These traits have not been explored in the F1 association mapping context, but we have previously demonstrated that *vey1* controls chlorophyll content in an F1 Association mapping experiment (Khangura *et al*. 2019).

We collected morphometric and time to reproductive maturity data from 343 F1 families that resulted from crosses between *Oy1-N1989*/+ in the B73 background and a densely genotyped maize diversity panel (**Figure 1**). Plant height and days to reproductive maturity were, on average, greater in the *Oy1-N1989*/+ F1 plants than in their wild-type siblings (**Figure 2**). The effects on plant height were primarily mediated by a change in the position of the ear and not by a change in the length of the stalk between the top ear and the flag leaf. By contrast, stalk width was consistently and dramatically decreased by *Oy1-N9189*. As expected, there was a substantial correlation between mutant and wildtype siblings for morphological traits (**Supplemental Table S2**) and to the chlorophyll content measurements performed on the same plants and reported previously (Khangura *et al*. 2019). To measure the impact of the mutant in each genetic background, we controlled for the contribution of trait variation in the isogenic wildtype siblings by calculating trait ratios (mutant/wildtype) or trait differences (wildtype-mutant) for each trait. Of the ten traits measured in mutants, only the anthesis-silking interval had a heritability lower than 0.80. Many heritability estimates for traits in the wildtype siblings, the ratiometric traits, and the WT-mutant difference traits also exceed 0.8 but were consistently lower than those measured in *Oy1-N1989*/+ mutant F1 families (**Table 1**).

**Figure 2.**
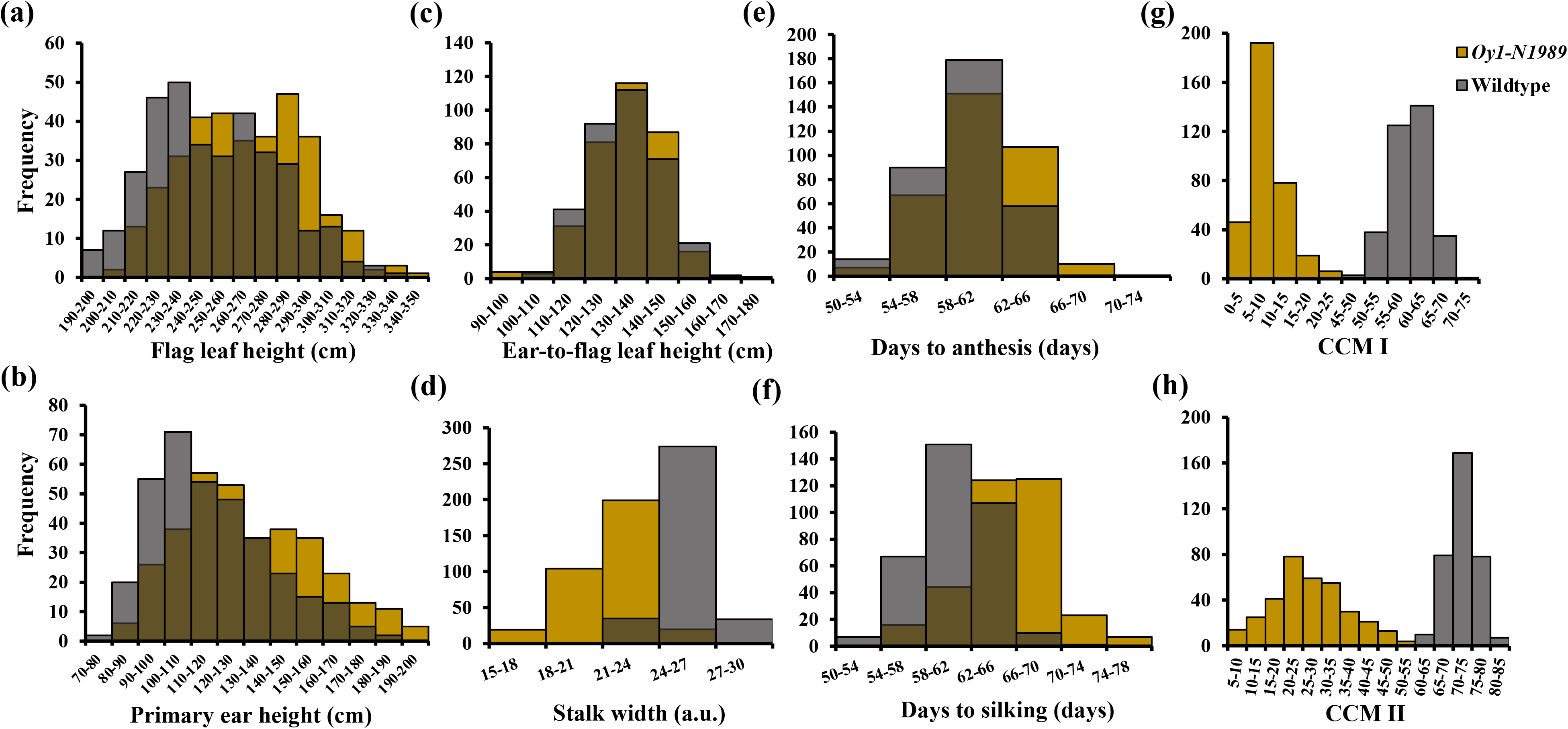
Frequency plot of the F1 hybrid wild-type and *Oy1-N1989*/+ mutant siblings. Overlaying trait distribution of isogenic *Oy1-N1989*/+ (golden color) and wildtype (grey color) siblings for (a) flag leaf height, (b) primary ear height, (c) ear-to-flag leaf height, (d) stalk width, (e) days to anthesis, (f) days to silking. Distributions from previously published dataset displaying (g) early season CCMI and (h) late season CCMII measurements in the same population.

**Table 1.** The descriptive statistics and broad-sense heritability (H^2^) of all traits measured in the F1 hybrids between the maize association panel and *Oy1-N1989*/+:B73 tester.

These phenotypic data, and the chlorophyll content measured previously (Khangura *et al*. 2019), were used to carry out genome-wide-association analyses. Consistent with the large effect of the allele-specific interaction between *vey1* and *Oy1-N1989*/+, all mutant traits, including chlorophyll content, stalk width, and days to reproductive maturity (both days to anthesis and silking), were modified by the QTL encoded at *oil yellow1* (**Figure 3; Supplemental Table S3**). SNP associations linked to *oy1* passed a Bonferroni corrected threshold that controls for genome-wide multiple tests. Controlling for the contribution of trait variation in the isogenic wildtype siblings by calculating trait ratios or trait differences did not erase the effect of the *vey1* QTL. GWAS using the difference and ratiometric values as traits detected top SNPs next to the *oy1* gene itself, except difference in flowering time (both differences in days to anthesis and differences in days to silking; diff_DTA and diff_DTS) where the lowest P-value SNP (10-9370704) was detected two genes away from *oy1* (**Figure 3b**).

**Figure 3:**
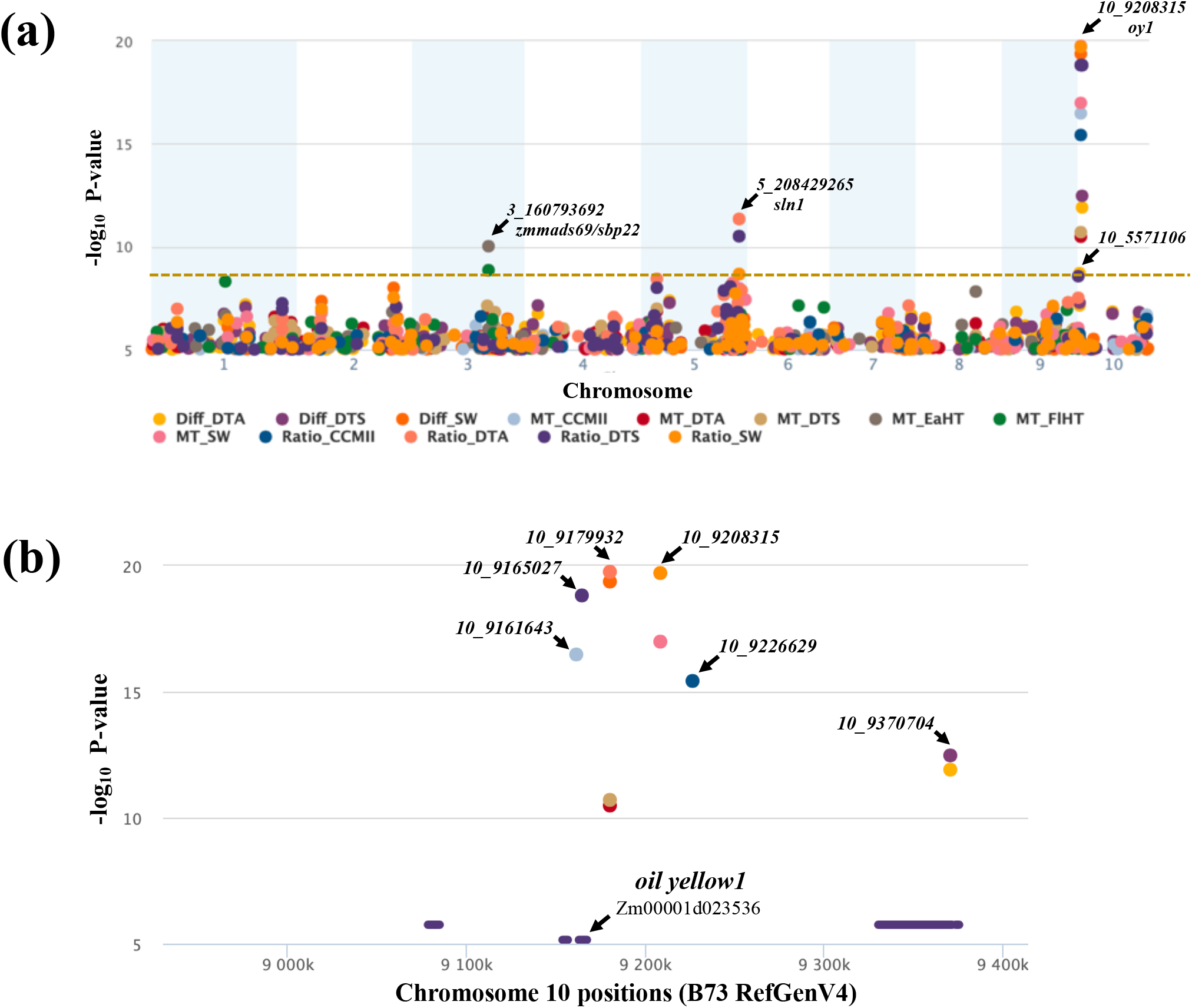
Manhattan plot showing the association test statistics in the MDLs X *Oy1-N1989/oy1*:B73 F_1_ test cross population. Golden hashed bar represents genome-wide bonferroni threshold. (a) Genome-wide plot of traits with at least one Bonferroni hit displaying only top SNP per trait in 250 kb region. (b) Close-up view of the significant SNPs at the *vey1* locus on chromosome 10.

In a single locus test using a B73 (with *vey1^Mo17^* introgression) and Mo17 (with *vey1^B73^* introgression) NIL population, we previously reported that *vey1*-dependent reduction of chlorophyll in *Oy1-N1989*/+ mutants resulted in thinner stems, delayed reproductive maturity, and reduced height. To test this relationship in the maize association panel F1 hybrids, we tested the top SNP associations for all traits linked to *oy1* for the effect direction of allele substitution on the trait. We used all six unique SNP-associations at *oy1* that had the lowest P-value for different mutant (including ratio and difference) traits and compared allele effect directions at these SNPs for all mutant and wild-type traits. All mutant traits (including ratio and difference) that had any SNP associations below a P-value of 0.0083 (Bonferroni cut-off for the test of six SNPs at alpha=0.05) showed a consistent effect direction where a decrease in chlorophyll conditioned by an allele also resulted in lower stalk width, delayed reproductive maturity, and reduced height. Consistent with no effect of *vey1* on the level of chlorophyll in the wildtype background, no consistent allele effects were observed in tests of wildtype traits (**Supplemental Table S4**).

Few loci other than *oy1* were detected at a Bonferroni corrected cut-off for any traits (**Figure 3; Supplemental Table S3**). One was located on chromosome 10, more than 3 Mbp away from *oy1*, at 5571106 bp (B73 RefGen v4), and affected the difference in DTA between wild type and *Oy1-N1989*/+ mutant hybrids (b=−2.4; p=1.9×10^−9^). This significant effect on the difference between wildtype and mutant DTA resulted from allele-specific interaction between SNP 10-5571106 and the genotype at *oy1* and no significant association was identified between this SNP and WT traits (**Supplemental Table S5**). DTA in the *Oy1-N1989* mutant and wildtype siblings were oppositely impacted by the alleles at 10-5571106 (*Oy1-N1989*/+: b=1.1, p=0.01; WT: b=−1.1; p=0.5). This locus may impact chlorophyll metabolism. A weak but significant effect was detected on chlorophyll content early in development in the mutants (CCMI; b=−3.8; p=0.006) and later for the CCMII ratio trait (b=−0.05; p=0.002). Consistent with previous findings (Khangura *et al*. 2020a), an alternate allele at this position decreased chlorophyll accumulation in the *Oy1-N1989* mutants and further delayed reproductive maturity. One candidate for QTL and association is a G-BOX BINDING TYPE B-ZIP transcription factor (Zm0001d023443), which is similar to the light-regulated gene expression determinants COMMON PLANT REGULATORY FACTOR2 (CFRP2) from parsley (Wellmer *et al*. 1999) and G-BOX BINDING FACTOR2 (GBF2), GBF3, BZIP16, and BZIP68 from Arabidopsis which are known to regulate nuclear-encoded photosynthetic components (Hsieh *et al*. 2012).

A flowering time QTL was detected on chromosome 5 at 208429465 bp (**Supplemental Tables S3 and S5**). This locus affected the ratio of mutant to wildtype DTA (b=0.03; p=4.5×10^−12^) and DTS (b=0.03; p=3.1×10^−11^). The alternate allele at this locus reduced mutant CCMII (b=−5.7; p=2 × 10^−4^) and increased mutant DTA (b=1.8; p=2 × 10^−4^) and mutant DTS (b=1.9; p=9.5×10^−5^). This is consistent with the observed relationship between chlorophyll accumulation and days to reproductive maturity. There are no known flowering time or chlorophyll regulators at this locus, but it is linked to the duplication of genes encoding delta-12-fatty acid desaturases (Zm00001d017840 and Zm00001d017841).

Another association was identified on chromosome 5 at 207300616 bp that affected both the ratio (b=−0.05; p=2.1×10^−9^) and difference (b=1.2; p=2.2×10^−9^) in stalk width between mutant and wildtype hybrid siblings (**Supplemental Tables S3 and S5**). At this SNP, an allele-specific interaction occurs where mutant stalk width was decreased (b=−1.0; p=2.4×10-7) and wildtype stalk width was increased (b=0.2; p=0.05) by the alternate allele. This locus also affected mutant CCMII (b=−3.2; p=7 × 10^−4^), which is consistent with the effect on the mutant stalk width being affected by a further decrease in chlorophyll accumulation. Consistent with a further reduction in chlorophyll by the alternative allele, mutant DTA was delayed (b=1.2; p=3 × 10^−4^). No effect was observed for plant height. No known regulators of chlorophyll metabolism or flowering time are linked to this position. However, the protein kinase *sister-of-liguleless narrow1* (Zm00001d029051; Buescher *et al*. 2014; Anderson *et al*. 2019) encodes a QTL that modifies the narrow stalk phenotype of *liguleless narrow1* (Zm00001d045945; Moon *et al*. 2013) is linked to this position and may explain this phenotype.

Multiple plant height QTL were detected for mutant height. Notably, associations that passed a Bonferroni-corrected threshold were located at loci in addition to *vey1/oy1*. Two SNPs on chromosome 3 passed a Bonferroni-corrected threshold affecting mutant ear height (position 160793692 bp; b=−7.9; p=9.5×10^−11^) and flag leaf height (160832709 bp; b=−7.9; p=1.4×10^−9^). The alternate allele at the 3-160793692 SNP accelerated reproductive maturity in the mutants for DTA (b=−0.9; p=1.5 × 10^−5^) and DTS (b=−0.8; p= 5.4×10^−5^). The alternate allele for the SNP at 3-160832709 bp also reduced reproductive maturity but the strength of the association was weaker (**Supplemental Tables S3 and S5**). These SNPs had no discernable effects on chlorophyll contents (p>0.05), indicating that these effects on growth and reproductive maturity were not mediated by altering chlorophyll accumulation in the impact of the *Oy1-N1989*/+ plants. As expected for *Oy1-N1989*-independent effects for SNPs influencing the same traits in the same direction as the wildtype siblings, we observed strong associations between the SNP at 3-160793692 bp and days to anthesis (b=−0.8; p=2.97 × 10^−7^), days to silking (b=−0.8; p=1.12x 10^−7^), ear height (b=−5.8; p=7.46 × 10^−8^), and plant height (b=−7.5; p=7.43 × 10^−8^) in the wildtype plants. For the SNP 3-160832709, similar to the mutant results, associations were weaker for wildtype days to anthesis (b= −0.5; p= 9.8 × 10^−4^), days to silking (b= −0.5; p= 6.8 × 10^−4^), ear height (b=−4.1; p=2.60 × 10^−5^), plant height (b=−6.3; p=8.15 × 10^−7^). We and others have previously identified a QTL linked to this position for wildtype plant height in B73 x Mo17 and teosinte x maize biparental populations (Liu *et al*. 2015; Liang *et al*. 2019; Khangura *et al*. 2020b). This position is linked to two known flowering time regulators, *zmmads69* (Zm00001d042315; Liang *et al*. 2019) and *zmsbp22* (Zm00001d042319; ~370 kbp upstream of *zmmads69*; Liu *et al*. 2020). The overexpression of either ZMMADS69 or ZMSBP22 is sufficient to accelerate flowering (Liang *et al*. 2019; Liu *et al*. 2020). ZMMADS69 also has pleiotropic effects on gross morphology, and overexpression of ZMMADS69 reduced ear height, plant height, and stalk width while accelerating flowering.

An alternative allele encoding an increased accumulation of ZMMADS69 or ZMSBP22 could underlie the flowering time QTL at this position. An eQTL analysis of RNA from the shoot apices of germinating seedlings from a B73 x Mo17 RIL population detected that the Mo17 allele increased the transcript abundance of ZMMADS69 but lowered the transcript accumulation of ZMSBP22 (**Supplemental Table S6**; Li *et al*. 2013, 2018). The Mo17 allele also decreased plant height and accelerated reproductive maturity in B73 x Mo17 populations (Liu *et al*. 2015; Khangura *et al*. 2020b). Thus, an eQTL at *zmmads69*, and not at *zmsbp22*, may encode the height and reproductive timing QTL at this position in the B73 x Mo17 cross. In our GWAS, we detected two closely-linked SNPs near *zmmads69* and *zmsbp22*, 3-160793692 and 3-160832709, affecting variation in plant height, ear height, and reproductive maturity in both mutants and wildtype siblings (**Supplemental Figure S1; Supplemental Table S7**). We carried out genetic associations between the 14294 SNPs in the ~1.4 Mb region containing ZMSBP22 and ZMMADS69 and their transcript count data from five green tissues (germinating shoot - GShoot, leaf three tip – L3Tip, leaf three base – L3Base, leaf at maturity during the day - LMAD, and leaf at maturity during the night - LMAN; Kremling *et al*. 2018) and compared this to the phenotypic data (**Supplemental Table S7**). Both SNPs were associated with increased expression of ZMMADS69 and were not significant in their effects on ZMSBP22. Thus, in both the GWA and RIL experiments, an eQTL explanation is only consistent with ZMMADS69 encoding the causative allele.

We detected a strong cis-acting eQTL for ZMMADS69 with 3-160559468 as the most significant SNP for transcript accumulation in multiple green tissues (Gshoot: p=7.3 × 10^−12^; b=0.4; **Supplemental Figure S1; Supplemental Table S7)**. The non-reference allele at this SNP increased ZMMADS69 accumulation and had consistent effects on wild-type and mutant sibling traits. This allele decreased plant height, ear height, and time to reproductive maturity (**Supplemental Table S7**). Thus, increased abundance of the ZMMADS69 transcript had the same effect on these phenotypes as the transgenic overexpression (Liang *et al*. 2019). At a threshold of p<10^−4^, only one tissue, L3Tip, contained any associations with ZMSBP22. None of the phenotypic traits were significantly affected by these SNPs. While they both nominally increased plant height, ear height, and reproductive maturity, one increased and the other decreased ZMSBP22 expression. Together with the weak effects on expression and lack of significant effects on plant growth traits, this rules out an eQTL at ZMSBP22 as a contributor to this allele.

In RIL experiments, the resolution of recombination is such that trait QTL and eQTL detection offer little opportunity to resolve linked effects of protein-coding versus gene expression differences. However, linkage in the maize association panel decays over much shorter physical distances, and recombinant genotypes with opposing effects on two traits can be detected (Remington *et al*. 2001). In the case of ZMMADS69 expression and the morphological QTL affecting flowering time and plant height, such recombinants are visible in the region. The top SNP (3-160559468) for ZMMADS69 accumulation results in a net decrease in days to reproductive maturity and a decrease in plant height in both mutant and wildtype plants (**Supplemental Figure S1; Supplemental Table S7**), consonant with the direction of effects observed in the RIL experiments (Khangura *et al*. 2020b). All 2198 SNPs that affected wildtype flag leaf height at p<0.001 resulted in a decrease in plant height and reproductive maturity and an increase in ZMMADS69 accumulation, demonstrating a linkage between trait and ZMMADS69 expression (**Supplemental Table S7**). Despite this linkage, ZMMADS69 accumulation is unlikely to be the cause of morphological trait variation. When the SNPs affecting ZMMADS69 accumulation in germinating shoots at a threshold of p<0.001 are examined (n=2156), the relationship between transcript accumulation and morphological effect is disrupted (**Supplemental Figure S2; Supplemental Figure S7**). A small number of SNPs that increased ZMMADS69 expression had a negative effect on plant height and hastened maturity. Thus, increased expression was associated with increasing and decreasing morphological trait values, indicating recombination between the causative polymorphisms responsible for ZMMADS69 expression and morphological variation. An even greater proportion of SNPs resulting in decreased ZMMADS69 accumulation were split into two groups that increased or decreased reproductive maturity and plant height. It is not possible for a reduced ZMMADS69 transcript abundance to both increase and decrease the population means of wildtype and mutant days to reproductive maturity and plant height. A similar decoupling of linkage between ZMMADS69 transcript accumulation and morphological traits was also observed using the L3Tip and L3Base expression data (**Supplemental Figures S2 and S3; Supplemental Table S7**). These data are consistent with at least two alleles linked in cis, with one allele affecting the expression of ZMMADS69 and another allele responsible for variation in days to reproductive maturity and plant height. As expected for this explanation, the SNPs that have opposing effects on transcript accumulation and plant growth do not disrupt the relationship between plant height and flowering time (**Supplemental Table S7**). The allele affecting the morphological traits underwent no recombination and is shared between these traits, while the allele affecting ZMMADS69 accumulation is not. Analysis of the protein-coding differences at ZMMADS69 in various maize inbred lines, including B73 and Mo17, or expression in the tissue responsible for the morphological effect, may resolve the molecular nature of this QTL.

### GWA using *vey1* as a covariate detected no additional loci

The frequent detection of allele-specific interactions between the GWAS QTL and our mutant and wildtype *oy1* alleles was striking. To further control for variation in chlorophyll affected at *oy1*, we modeled the effect of *vey1* and scanned the remainder of the genome for genes influencing the *Oy1-N1989*/+ hybrid phenotypes. To do this, we used the top SNP associations at *vey1* as covariates in a GWAS to control for the variation affected by these large-effect QTL. We iteratively ran GWAS for mutant CCMII and added the top SNP linked to *oy1* as a covariate until no SNP linked to *oy1* exceeded a p-value of 10^−6^ for this trait. This resulted in a model with three SNPs (10-9161643, 10-9179932, and 10-9226629) that impacted mutant CCMII. We included these three SNPs as covariates in the model to run GWA for all other traits. The only covariate-corrected GWA locus that passed the genome-wide Bonferroni threshold for any trait was the previously detected QTL for mutant ear height and mutant flag leaf height linked to *zmmads69* and *zmsbp22* (**Supplemental Table S8**). This result suggests that the allelic difference linked to *zmmads69*/*zmsbp22* did not alter morphology via a change in chlorophyll content by the *Oy1-N1989* allele. Thus, controlling for the large effect at *vey1* did not uncover any additional QTL for chlorophyll content, reproductive maturity, or plant height.

### Evaluation of genes in tetrapyrrole biosynthesis pathway as candidates for *Oy1-N1989* suppression

The *oy1* gene encodes the Mg-Chelatase subunit I, and variation in other genes involved in tetrapyrrole metabolism might enhance or suppress the impact of this mutant (**Figure 4**). Besides the *vey1* association at *oy1* itself, no genes of known function in biosynthesis or regulation of tetrapyrrole were among the loci linked to a strict genome-wide Bonferroni threshold. Therefore, we carried out candidate-gene associations with this pathway to determine if allelic variation in tetrapyrrole metabolism genes explains any variation in chlorophyll accumulation in the *Oy1-N1989* diversity population. We tabulated all the SNP associations with a P-value of less than 10^−4^ and retained the lowest P-value SNP (also referred to as a “*top SNP*”) within a 250kb window. Consistent with additional loci affecting chlorophyll content in the mutants, there were more loci at this threshold for mutant chlorophyll accumulation (750 for CCMI and 410 for CCMII) than for the wildtype siblings (268 for CMMI and CCMII 321). The difference and ratio traits, which control for the baseline trait in the wildtype siblings, detected 297 top SNPs for difference CCMI, 424 for difference CCMII, 521 for ratio CCMI, and 460 for ratio CCMII. The combination of these SNP-trait associations resulted in 2522 unique SNPs, with some in close proximity to each other. There are 57 genes in the maize genome known to encode enzymes in the biosynthesis and degradation of chlorophyll, heme, and other porphyrins (**Supplemental Table S9**). This list of genes was compared to the set of 2522 top SNPs, and any *top SNP* within three genes of a porphyrin gene was retained as a possible candidate association. There were 14 genes (including *oy1*) that were tagged by a SNP with a significant association at P-value of 10^−4^ or less for at least one CCM trait. The effect of these alleles on plant height, stalk width, and reproductive maturity traits were also tabulated for comparison (**Supplemental Table S10**).

**Figure 4.**
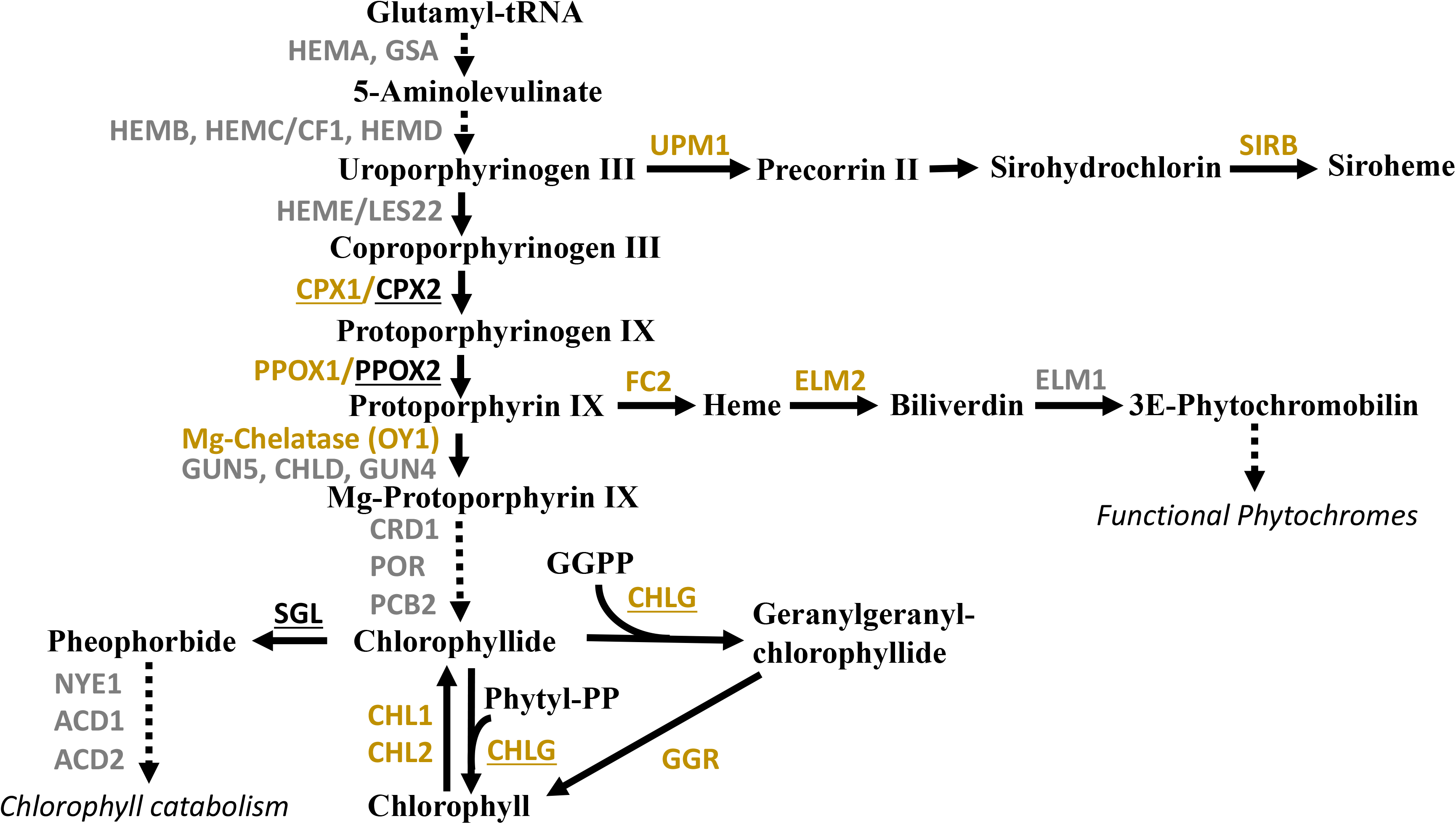
Schematic of porphyrin, chlorophyll, heme, and siroheme biosynthesis pathways. Enzymes with linked SNPs with associations at p-value < 10^−4^ with chlorophyll traits in *Oy1-N1989*/+ F1 families are golden. Enzymes linked to SNPs with associations at p-value < 10^−4^ with chlorophyll traits in WT sibling families are underlined. Other enzymes with no significant SNP associations are shown in grey. Data in **Supplemental Table S8**.

We identified alleles linked to enzymes in porphyrin, heme, siroheme, and chlorophyll metabolism with significant effects on chlorophyll variation in the mutant or mutant and wildtype comparisons (**Figure 4**). In the porphyrin biosynthesis pathway, we identified SNP associations of mutant CCM with COPROPORPHYRINOGEN III OXIDASE I (CPX1; Zm00001d002358), COPROPORPHYRINOGEN III OXIDASE II (CPX2; Zm00001d026277), and PROTOPORPHYRINOGEN IX OXIDASE I (PPOX1; Zm00001d008203). A paralog of PPOX1, PROTOPORPHYRINOGEN IX OXIDASEII (PPOX2; Zm00001d003214), was only associated with wildtype CCM values. FERROCHELATASE II (FC2; Zm00001d016854), and HEME OXYGENASE I/ELONGATED MESOCOTYL 2 (HO1/ELM2; Zm00001d046492), which are involved in heme and bilin biosynthesis, were associated with mutant CCM values. In the siroheme pathway, which is necessary for nitrogen and sulfur metabolism, SIROHEME UROPORPHYRINOGEN METHYLTRANSFERASE I (SUM1 or UPM1; Zm00001d044138) and SIROHYDROCHLORIN FERROCHELATASE B (SIRB; Zm00001d002511) were associated with mutant CCM values. In chlorophyll metabolism, CHLOROPHYLLASE I (CHL1; Zm00001d019758), CHLOROPHYLLASE II (CHL2; Zm00001d032926), CHLOROPHYLL SYNTHASE G1 (CHLG1; Zm00001d037984), STAY GREEN-LIKE (SGL; Zm00001d002951) were associated with CCM. Consistent with a demonstrated effect on leaf greenness (Shimoda *et al*. 2016), SGL only influenced wildtype CCM values. Loci encoding CHLG1 and GERANYLGERANYL REDUCTASE (GGR; Zm00001d018034) that are required for utilizing the phytyl side chain from geranylgeranyl diphosphate were associated with mutant CCM values. These alleles associated with mutant CCM also affected flowering time and stalk width in the expected direction, where low CCM alleles resulted in plants with thinner stems and delayed reproductive maturity. A complete description of the genes in **Figure 4** and the effects of the alleles are presented as supplementary text **(Supplementary text file S1)**.

### Associations with *bona fide* flowering time regulators

We tested whether SNP-to-trait associations for reproductive maturity and height traits were linked to known regulators of flowering time in maize. Due to a known pleiotropic effect of flowering time regulators on height in maize (Danilevskaya *et al*. 2011), we searched for alleles at these genes for both sets of traits. Again, we tabulated all the SNP associations with a P-value of less than 10^−4^ for these traits and retained the lowest P-value SNP within a 250kb window. A set of 18 *bona fide* flowering time genes in maize (**Supplemental Table S11**) were compared to the set of three genes up and three genes down from the top SNP for each flowering time and plant height locus. Thirteen of the eighteen genes (72%) encoding *bona fide* flowering time regulators (*zmsbp22*, *zmmads69*, *id1*, *d8*, *d9*, *zfl2*, *dlf1*, *zcn8*, *zmtps1*, *zmrap2*, *conz1*, *zmm1*, and *cct1*) were within three genes of one of 29 top SNPs defining a reproductive maturity or height traits locus in either mutant or wildtype siblings (**Supplemental Table S11**). Because of their proximity, *zmsbp22* and *zmmads69* were linked to seven SNPs. None of the 29 SNPs impacted CCM in either mutant or wildtype siblings at the P-value threshold of 10^−4^ used for this analysis or after correcting for 29 multiple tests (Bonferroni alpha<0.05 equivalent to p< 0.0017; **Supplemental Table S13**). This indicates that these loci were not influencing reproductive maturity or height in *Oy1-N1989*/+ F1 plants via an indirect effect on chlorophyll, as observed for *vey1*. A subset of SNPs linked to *zmsbp22*/*zmmads69, d9*, *d8*, and *zfl2* all had effects on CCMII mutant to wildtype ratio with uncorrected p<0.05 (**Supplemental Table S12**). Other than *zmsbp22*/*zmmads69*, which passed a Bonferroni threshold as discussed above, we further investigated the associations with the remaining eleven *bona fide* FT regulators. A detail on the effects at these SNPs on flowering time regulators are provided in the supplemental text **(Supplementary text file S1)**, but its synthesis is summarized below.

Where a SNP-to-trait association was significant in either wildtype or mutant sibling populations, the direction of the effect was always the same in both populations. This contrasts with the allele-specific effects observed for the GWAS QTL that influenced flowering time and stalk width via chlorophyll (above) that either had opposite effects on mutant and wildtype siblings or only affected phenotypes in mutant plants. This suggests that the delay in flowering affected by *Oy1-N1989*/+ does not interact with pathways specified by these known flowering time loci. Even in cases where the traits were calculated as the mutant to wild type ratio, the majority were affected in the same direction in both. The exceptions were a small number of SNPs linked to *d8* and *zmm1* for height traits and *d9* and *zfl2* for DTA and DTS (**Supplemental Table S12**). These genes encode alleles with nominally significant effects on the CCMII ratio. However, the effects on chlorophyll were unlikely to cause the change in morphology as the effect directions were inconsistent. Alleles at *d8* and *d9* influenced the ratio of mutant to wildtype heights suggesting that GA may be required for *Oy1-N1989* to modify plant height.

### GWA as a fine-mapping strategy for previous QTL studies using *Oy1-N1989*/+ tester

We previously identified multiple loci that modify the *Oy1-N1989*/+ mutant by QTL mapping in F1 crosses to B73 x Mo17 RIL. Biparental mapping populations between B73 and Mo17 were crossed to the *Oy1-N1989*/+ mutant tester in B73 inbred background as a pollen-parent (Khangura *et al*. 2019, 2020a; b), and the F1 progeny were evaluated for a variety of traits. 42 QTLs were detected, 23 of which were encoded by a large-effect QTL at *oy1*, which we called *vey1*. The 19 additional loci detected in these experiments affected variation in height, days to reproductive maturity, and stalk width in either mutant or wildtype siblings (Khangura *et al*. 2020a; b). We determined if the 2-LOD genetic intervals of these 19 QTL loci colocalized with any GWAS associations. As we considered a 2-LOD window for these QTL, rather than genome-wide, we carried out these candidate associations at a threshold of P-value=10^−4^. Of the 2-LOD intervals in these 19 QTLs, 8 contained at least one SNP with a P-value below our threshold for any trait. These 8 QTLs are encoded at five distinct loci, with three loci affecting two traits each (**Supplemental Table S13**). We calculated the effect of the non-B73 (non-reference or alternate) allele so that we could directly compare the effect direction of any GWAS result to our prior B73xMo17 QTL results. A detailed comparison of GWAS and QTL allele effect direction and candidate genes were performed within the narrower association window for these five loci.

Two QTL, one affecting DTA in mutants and the other DTS in mutants, resulted from the same underlying allele, which we previously named *other oil yellow flowering locus1* (*oof1*; 2-LOD chr2: 96.1-103.4 Mbp; Khangura et al., 2020a). The SNP with the lowest P-value in this 2-LOD window for reproductive maturity in mutants is at chr2-102476588 bp, and the alternate allele delays both mutant DTA (P=9.4 × 10^−5^; b= −1.28) and mutant DTS (P=1.3 × 10^−4^; b= −1.28). Although this QTL was identified for mutant reproductive maturity, a modest association was also observed at this SNP that also delayed both wildtype DTA (P=3.2 × 10^−4^; b= −0.9) and DTS (P=2.5 × 10^−4^; b= −0.9; **Supplemental Table S14**). No known regulators of flowering time in maize were present at this position. The hastened maturity by the alternate allele at this SNP was dissonant with our previous study, where a delay was observed with the Mo17 allele (**Supplemental Table S13**). Thus, it is unlikely that this GWAS association is responsible for the *oof1* QTL.

For height traits in mutants, we previously detected a total of four QTL. Only one of these QTLs overlapped with a significant SNP association. This QTL was identified in B73 x Mo17 at chr1-622 cM (2-LOD chr1: 82.3-82.5 Mbp) and affected both mutant and wild-type sibling heights. This QTL overlapped a SNP (chr1 at 82335623 bp) in our association mapping experiment that was associated with the alternate allele increasing mutant ear height (P=7.2 × 10^−5^; b= 5.7) and wildtype ear height (P=4.8 × 10^−4^; b= 4.5). A modest association of this position was observed with flag-leaf height for both mutant and wild-type siblings. The effect direction of the alternate allele at this SNP was consonant for both genotypes and was also consistent with the direction of the Mo17 allele described previously (**Supplemental Table S13**). This SNP is linked to a MATE transporter (Zm00001d029677) that we previously proposed as a candidate gene for this QTL (Khangura *et al*. 2020b).

For height traits in the wildtype siblings, we previously identified a QTL on chromosome 1 for wildtype ear-to-flag-leaf (Ea2Fl) height and another QTL for wildtype flag-leaf (FlHT) height at overlapping genomic intervals, with the Mo17 allele resulting in a reduction of plant height (**Supplemental Table S14**). The lowest P-value SNP in this interval was chr1-189266249 bp, and the alternate allele decreased wildtype Ea2Fl (P=4.1 × 10^−5^; b= −2.5) and to a lesser extent reduced wildtype FlHT (P=0.01; b= −3.4). No gene with a known impact on plant height was identified at this position. On chromosome 3, we previously reported a QTL (2-LOD chr3:160.6-160.7 Mbp) that co-localized with *zmmads69* (Zm00001d042315) and *zmsbp22* (Zm00001d042319), both of which are known flowering time and height regulators in maize (Liang *et al*. 2019; Liu *et al*. 2020). Just like the Mo17 allele at this QTL that we described previously, in this study, a SNP at chr3-160607046 bp decreased ear height (mutant: P=1.5 × 10^−7^; b= −5.8, wildtype: P=3.7 × 10^−5^; b= −4.1), flag-leaf height (mutant: P=1.3 × 10^−6^; b= −6.1, wildtype: P=1.5 × 10^−5^; b= −5.5), and resulted in hastening of reproductive maturity (mutant DTA: P=2.8 × 10^−5^; b= −0.8, wildtype DTA: P=1.8 × 10^−5^; b= −0.6) in both mutant and wildtype siblings.

Lastly, for stalk width, only one QTL from our previous study, on chromosome 1 (2-LOD chr1: 254.4-255.9 Mbp), colocalized with a SNP association in this study. Like the Mo17 allele from our previous work, the non-B73 allele at this SNP (chr1 at 55224320 bp) increased wildtype stalk width (P=3.7×10^−5^; b=0.3). This SNP was linked to a *β-expansin4 (expb4*; Zm00001d033231), a maize gene we proposed as a candidate gene in our previous QTL study (Khangura *et al*. 2020b).

### Reproductive maturity delay in *Oy1-N1989* is due to a delay in growth and not phase transition

The delay in reproductive maturity observed in *Oy1-N1989*/+ could be due to a delay in growth rate or a developmental delay. The timing of juvenile to adult phase change, rate of new leaf production, and the number of phytomers between stages were measured to distinguish between these two mechanisms. The number of leaves required for the juvenile to adult phase change was unaltered in *Oy1-N1989*/+, and wild-type siblings produced a fully adult leaf by leaf 8 (**Table 2**). Enhancement of the *Oy1-N1989*/+ phenotype by the *vey1^Mo17^* QTL did not delay phase change either (**Table 2**), despite these plants reaching reproductive maturity approximately two weeks later than wildtype plants (Khangura *et al*. 2020a). Similarly, the total number of leaves was not affected by *Oy1-N1989*/+ or its enhancement by the *vey1^Mo17^* allele (**Table 3**). Other attributes of plant development, such as leaves below and above the primary ear, were also not affected (**Table 3**). The leaf emergence rate after the phase change was dramatically reduced in *Oy1-N1989* mutants compared to the wildtype siblings in both B73 inbred and Mo17 x B73 hybrid combinations (**Figure 5**). Enhancement and suppression of *Oy1-N1989* mutants by the *vey1^Mo17^* and *vey1^B73^* alleles slowed and accelerated leaf production, respectively (**Figure 5**). This demonstrates that the delay in reproductive maturity was affected by *Oy1-N1989*, and enhancement of this delay by the *vey1^Mo17^* allele resulted from slower growth and not an alteration in the developmental program.

**Table 2.** The juvenile-to-adult leaf transition determined by changes in epicuticular leaf wax in wildtype and *Oy1-N1989*/+ mutant siblings in B73 inbred and Mo17 x B73 F1 hybrid backgrounds along with reciprocal introgression of *vey1* alleles.

**Table 3.** The number of leaves are not affected due to *vey1* x *Oy1-N1989* interaction.

**Figure 5.**
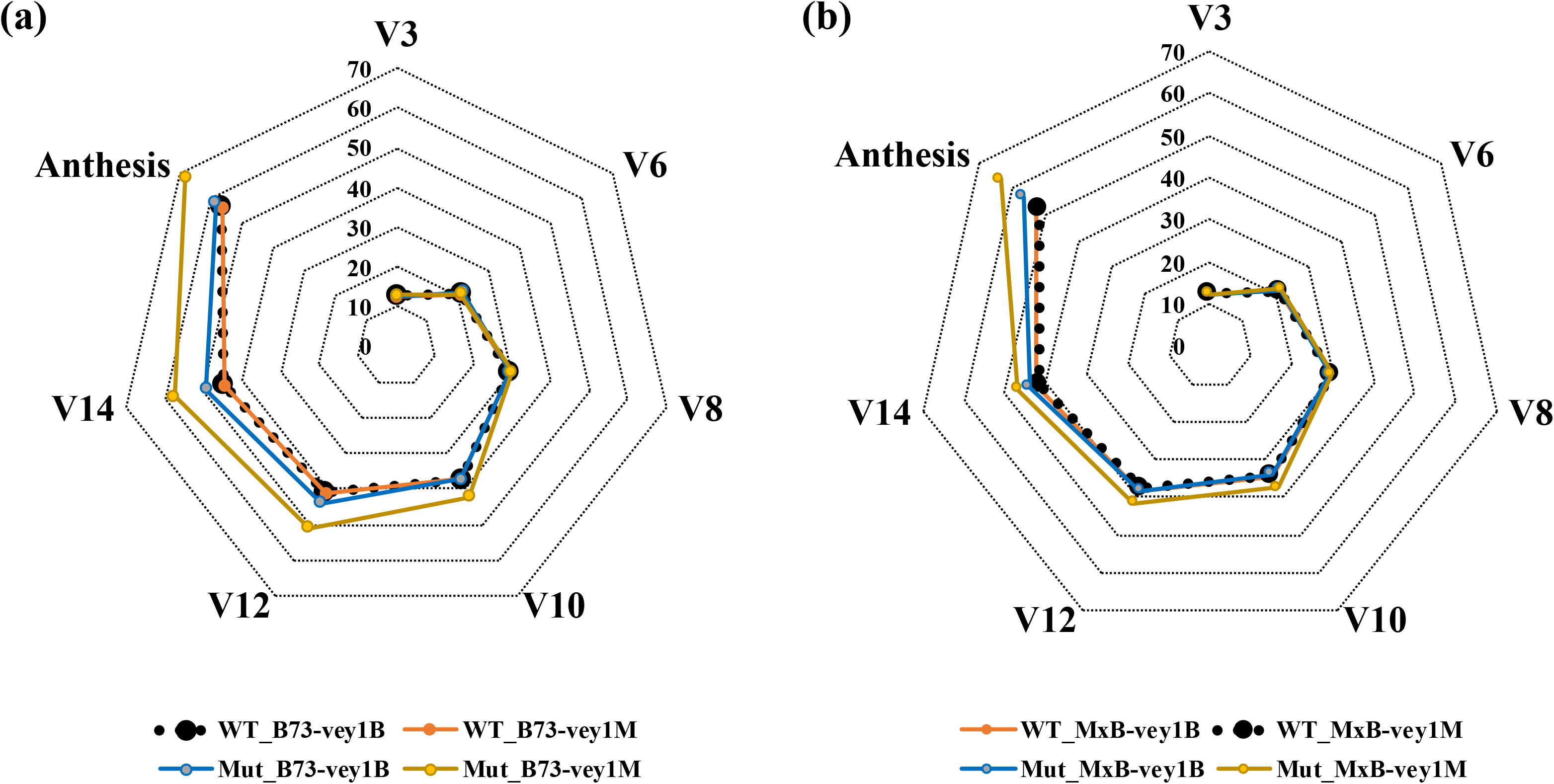
Rate of leaf emergence and anthesis in the wildtype and *Oy1-N1989*/+ siblings in isogenic genetic backgrounds (A) B73, and (B) Mo17 x B73 F1 hybrids. Abbreviations - WT: wildtype, Mut: *Oy1-N1989*/+ mutant, B: B73, M: Mo17.

### Relationship of CCM and *Oy1-N1989*/+ suppression to height

We previously observed a complex relationship between CCM and plant height in *Oy1-N1989*/+ plants. Crosses between *Oy1-N1*989 and the B73 x Mo17 RILs produced two classes of *Oy1-N1989* mutants: those enhanced by *vey1^Mo17^* and those suppressed by *vey1^B73^*. All mutants enhanced by *vey1^Mo17^* were shorter than wild-type siblings and had CCMII values below 9 (Khangura *et al*. 2020b). Further declines in CCMII were associated with a reduction in plant height. All mutants suppressed by *vey1^B73^* had less chlorophyll than wildtype plants but had CCMII values above 11 and were taller than their wildtype siblings (Khangura *et al*. 2020b). Further declines in CCMII values in these plants did not affect plant height, and they remained taller than their wildtype siblings despite being yellow-green (Khangura *et al*. 2020b). No known mechanism can explain how the loss of chlorophyll can modify plant height in opposing directions. We proposed that there is a positive height impact of reducing chlorophyll that is eventually offset when photosynthesis is sufficiently limited, and this impedes growth. Therefore, there is likely an optimal chlorophyll concentration at which mutant plant height is maximized. Unlike the crosses to B73 x Mo17 RILs, the CCMII values in the F1 association mapping populations varied from 61.4 – 82.4 in the wildtype F1 hybrids and displayed a continuous range from 5.9-53.5 in the *Oy1-N1989*/+ mutant F1 hybrids (**Figure 6**). This uninterrupted distribution of chlorophyll levels in the mutants allowed us to determine the chlorophyll level that maximized plant height of *Oy1-N1989*/+ mutants with respect to its wild-type siblings.

**Figure 6.**
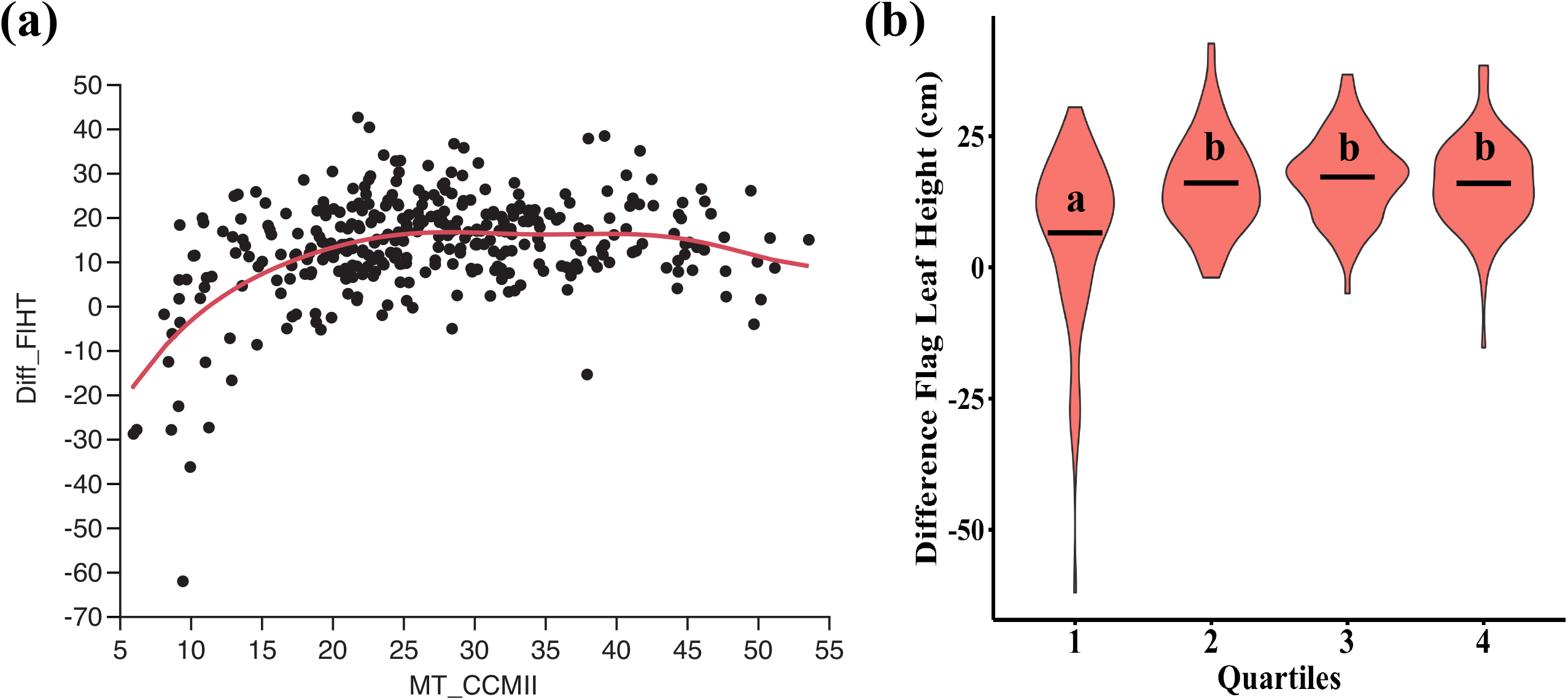
Non-linear relationship of mutant CCM with flag leaf height difference between isogenic F1 mutant and wildtype siblings. (a) Lowess curve fit showing deviation from a linear fit. (b) Distribution of difference in flag leaf height in each quartile (as determined by mutant CCMII). The black bars indicates mean, and the connecting letter report indicates statistical significance between means determined using student’s t-test at p<0.05.

In the *Oy1-N1989*/+ mutant F1 hybrids, CCMII and plant height displayed a non-linear relationship with an optimum chlorophyll level at which height was maximized (**Figure 6**). A smoothed LOESS fit of the data confirmed a non-linear relationship between height difference and mutant CCMII at both ends of the chlorophyll distribution (**Figure 6**). A CCMII of 21.8 coincided with the peak of the linear fit and corresponded to an increase in mutant plant height of 42.7 cm above its wild-type siblings. Similar to our previous observations in RILs, at mutant CCMII values less than 15, the *Oy1-N1989*/+ mutants were shorter than their wild-type siblings (Khangura *et al*. 2020b). To further explore the non-linear relationship between height difference and mutant CCMII, we divided the dataset into quartiles using mutant CCMII values (**Figure 6**). Analysis of quartiles showed a similar trend where the mutants with the most severe reductions in CCMII were shorter than the other three quartiles. The last three quartiles displayed a sustained increase in mutant plant height over their wild-type siblings. These data are consistent with an increase in plant height in *Oy1-N1989*/+ induced by moderate reductions of chlorophyll content starting just below the wildtype range and progressing to ~ 22 units of mutant CCMII. A further loss of chlorophyll below 15 units of CCMII resulted in a progressive and dramatic decrease in mutant plant height compared to its wild-type siblings (**Figure 6**). Thus, a “goldilocks” model for controlling plant height by chlorophyll content modulation proposed in our previous work using B73 x Mo17 population (Khangura *et al*. 2020b) was confirmed in the F1 association mapping population. While limitations to the photosynthetic rate at lower CCM values in *Oy1-N1989*/+ mutants (Khangura *et al*. 2020a) might limit plant growth, there is no known mechanism for chlorophyll limitation to result in taller plants. However, phytochromes and other porphyrin derivatives are known to influence plant height.

### *elongated mesocotyl1* and *Oy1-N1989* double mutants are synthetically deleterious

The Mg-Chelatase subunit I encoded by *oy1* uses protoporphyrin IX, the last shared precursor of the heme and chlorophyll biosynthetic pathways, as a substrate (**Figure 4**). 3E-phytochromobilin is a heme derivative that is the chromophore of phytochrome. In maize, mutations in phytochromobilin synthase encoded by *elongated mesocotyl1* (*elm1*; Zm00001d011876) disrupt chromophore production and result in pale green plants that are taller than the wildtype siblings, a phenotype reminiscent of the tall suppressed *Oy1-N1989*/+ mutants (Sawers *et al*. 2002, 2004). We constructed double mutants to test the interaction between *elm1-ref* and *Oy1-N1989*. The *elm1-ref* single mutants had a CCMI of 18.5, The *Oy1-N1989*/+ single mutants had a CCMI of 4.8, and the wildtype plants had a CCMI of 29.6. Thus *elm1-ref* and *Oy1-1989*/+ had two- and six-fold less chlorophyll than the wildtype siblings, respectively (**Figure 7**). The double mutants between *elm1-ref* and *Oy1-N1989* were additive for their effects on chlorophyll content resulting in a CCM value of 2.2 (**Figure 7; Supplemental Table S16**). The *Oy1-N1989/+; elm1-ref* double mutants had six-fold less chlorophyll than wild-type siblings (**Figure 7**). Only *Oy1-N1989*/+ single mutants, but not *elm1-ref*, were significantly taller than their wild-type siblings. The *elm1-ref;Oy1-N1989*/+ double mutants, however, demonstrated a synthetically deleterious effect on plant height resulting in plants shorter than wildtype siblings by 56 cm. The *elm1-ref* and *Oy1-N1989*/+ single mutants have opposite effects on the timing of reproductive maturity, with *elm1-ref* accelerating and *Oy1-N1989* delaying reproduction (**Figure 7**; and Sawers *et al*. 2002; Khangura *et al*. 2020a). The double mutants resembled a strongly enhanced *Oy1-N1989*/+ mutant, and their anthesis was delayed by 13 days (**Figure 7**). Both *elm1-ref* and *Oy1-N1989* mutants decreased stalk width. The double mutant phenotype was strongly synergistic, resulting in a greater-than-additive decrease in stalk widths in the *Oy1-N1989*/+;*elm1-ref* double mutants. For all three phenotypes, the non-additive synthetic effects of *elm1-ref* and *Oy1-N1989*/+ on plant growth suggest that they are controlling growth as part of a shared pathway. This implicates feedback regulation of porphyrin metabolism as a mechanism for reducing the CCM values of *elm1-ref* mutants and synergistic interaction between *Oy1-N1989*/+ and *elm1-ref*.

**Figure 7.**
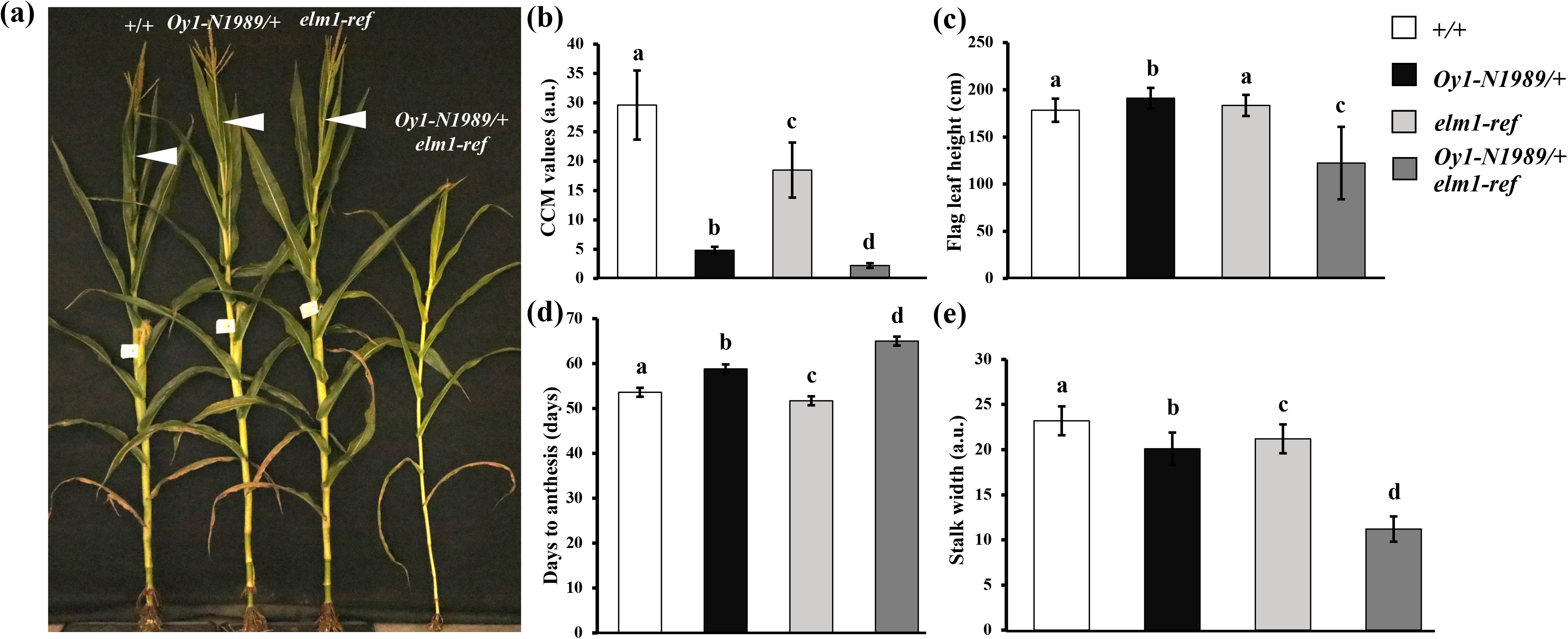
The single and double mutant analysis between *Oy1-N1989*/+ and *elm1-ref* in B73 inbred background. (a) Photograph of a representative wild-type (+/+), *Oy1-N1989*/+;*elm1-ref*/+, *elm1-ref/elm1-ref*, and a *Oy1-N1989*/+;*elm1-ref/elm1-ref* individuals from a segregating F2 family in B73 background at maturity. Average trait distribution of all four genotypes for (b) CCM, (c) flag leaf height, (d) days to anthesis, and (e) stalk width. Error bars indicate standard deviation with connecting letter denoting significant difference between means at p<0.05 as determined by student’s t-test.

## Discussion

We previously identified a major cis-eQTL at *oy1*, which we call *vey1*, that is predominantly responsible for the variation in *Oy1-N9189*/+ phenotype expression between B73 and Mo17 (Khangura *et al*. 2019, 2020a; b). This eQTL, and detailed study of mutant expression in RIL and NIL F1 *Oy1-N1989*/+ plants with contrasting *vey1* alleles, determined that time to reproductive maturity, plant height, and stalk width were affected by chlorophyll content in maize (Khangura *et al*. 2019, 2020a; b). A GWAS of chlorophyll content in *Oy1-N1989*/+ mutants also mapped a modifier acting in cis, further indicating that *vey1* was encoded by an allele of *oy1* (Khangura *et al*. 2019). The current study further explores the relationship between chlorophyll and plant morphology in maize.

### Candidate genes from pathway-level analyses

Because the mapping resolution of GWA allows a finer-scale exploration of candidate genes, we returned to those proposed in our previous work. We performed pathway-level analyses to determine if alleles in genes in the tetrapyrrole pathway affected variation in chlorophyll levels in *Oy1-N1989*/+ plants (**Figure 4**). This study reproduces our previous observations and addresses some of the questions raised in our previous work. A genome-wide Bonferroni threshold will result in a high rate of false negative assessments. The general bias of the field is to not encourage the expensive and time-consuming work of single locus mapping or gene validation without strong evidence for a candidate gene for natural variation. When the pathway is known, a search for alleles at a subset of loci can still be corrected for genome-wide multiple-test corrections, but the number of possible SNPs tested is lower. Because we know that Mg Chelatase in the porphyrin biosynthetic pathway, focusing on other genes known to encode steps in this pathway returned multiple alleles likely to encode weak modifiers of small effect. In fact, out of 57 genes broadly defined as in the tetrapyrrole pathway in pathway, 11 were linked to SNPs with a P-value less than 10^−4^ affecting mutant chlorophyll content, and only four affected wildtype CCM (**Figure 4; Supplemental Table S9)**. This fits with the observation that even *vey1* has no effect on chlorophyll content in the wild-type siblings and suggests that the search for modifiers of *Oy1-N1989* has uncovered additional cryptic variants for which *Oy1-N1989* is epistatic.

We reanalyzed our previously proposed candidate genes from the line cross populations (Khangura *et al*. 2020a; b). In some cases, the narrower window provided by GWAS still contained the candidates we previously proposed, such as *zmmads69*, the *expansin* encoded by Zm00001d033231, and a mate transporter Zm00001d029677. In two other cases, we previously did not propose candidates during the analysis of our line-cross QTL.

### How does *Oy1-N1989* impact flowering time?

We previously demonstrated that *Oy1-N1989*/+ mutants had a reduced ability to assimilate CO_2_ (Khangura *et al*. 2020a). We showed that reduction in CO_2_ assimilation also reduced levels of non-structural carbohydrates, namely starch, sucrose, glucose, and fructose. This reduction in carbon assimilation and sugars was sensitive to the *vey1* allele, where a severe reduction in chlorophyll by the *vey1^Mo17^* allele results in greater reductions than the suppressing *vey1^B73^* allele (Khangura *et al*. 2020a). Carbon sensing has been proposed to play an important role in controlling flowering time. In maize, INDETERMINATE1 and Hexokinases were proposed to regulate the reproductive transition (Minow *et al*. 2018). In addition, sugar is proposed to promote vegetative to adult phase change via direct modulation of miR156a and miR156c (Yang *et al*. 2013). An alternative possibility is that source limitation results in slower growth and does not delay development. We demonstrate here that the *Oy1-N1989* allele does not impact the juvenile-to-adult phase transition and the number of phytomers, nor is it altered by a change in chlorophyll levels due to the *vey1* modifier (**Tables 2 and 3**). Instead, the *Oy1-N1989*/+ mutants exhibit slower leaf emergence from the whorl than \ wildtype siblings, and this is more severe when enhanced by *vey1^Mo17^* (**Figure 5**). This effect is visible in both isogenic B73 and Mo17 x B73 F1 hybrid backgrounds. Thus, leaf emergence is delayed in the *Oy1-N1989*/+ mutants due to the reduced level of photosynthates in a *vey1*-dependent manner. We propose that the delay in reproductive maturity in the *Oy1-N1989*/+ mutants is due to a slower growth rate and not a developmental delay at the meristem.

All known flowering time regulators with observed effects in our F1 mapping experiments had a similar impact on both the wildtype and the mutant siblings (**Supplemental Table S13**). This provides further evidence that the reproductive delay was affected by the reduced growth rate in *Oy1-N1989*/+ and not any known regulators of flowering time. It also demonstrates that a dramatic reduction in photosynthesis-derived carbon did not affect flowering time, as the number of phytomers was unchanged. This runs counter to the proposal of the role of a carbon sensor in regulating the autonomous pathway of flowering time in maize (Minow *et al*. 2018)

### What makes *Oy1-N1989*/+ plants tall?

The *Oy1-N9189*/+ mutants were taller than the wildtype siblings when the chlorophyll phenotype of the mutant was weak, and the mutants got progressively taller as the chlorophyll contents decreased further (**Figure 6**). However, strong enhancement of *Oy1-N1989*/+ resulted in shorter plants, and mutant plants got progressively shorter with a further decline in chlorophyll contents. Thus, a decrease in chlorophyll could have a positive or a negative effect on plant height. This explains why we did not observe a strong QTL at *vey1* for the mutant height traits (**Figure 3 and Supplemental Table S3**). Despite the demonstrated effects of *Oy1-N1989* and the *vey1* modifier on plant height, no associations were seen for plant height effects at *vey1* (**Figure 3; Supplemental Table S3**). If we limit our view to just the SNP with the strongest effect on CCM at *vey1* (10-9161643), the CCM-decreasing alternate allele has a weak negative effect on mutant plant height (p=0.006; b=−3.4). We have now observed this phenomenon in two biparental mapping populations (Khangura et al., 2020a), NILs (Khangura et al., 2020a), and our F1 diversity panel (**Figure 3; Supplemental Table S3**). The reduction of plant height by a severe decline in chlorophyll levels is no surprise. Plants defective in photosynthesis lack the fixed carbon to support growth. However, we need to find a molecular explanation for why partially suppressed *Oy1-N1989* mutants are taller.

Partially suppressed *Oy1-N1989*/+; *vey1^B73^* plants have 15-30% lower net CO_2_ assimilation rates (Khangura *et al*. 2020a). This is sufficient to delay days to reproductive maturity and decrease stalk width, demonstrating that these reductions in carbon assimilation are sufficient to affect phenotypes. Yet, these mutant plants are taller than wild-type siblings (**Figure 7** and Khangura *et al*. 2020b). Regardless of the mechanism for the increase in height, we expect that as chlorophyll levels fall, plants get smaller. This is what is observed for stalk width. The presence of some yet unknown mechanism for *Oy1-N1989* to increase plant height results in modest increases in height until the negative consequence of poor photosynthesis overwhelms the growth capacity of the plant, resulting in shorter plants when the phenotype is more severe. One attractive hypothesis we have explored is that the effects of *Oy1-N1989* on the porphyrin pathway result in a change in bilin and phytochrome signaling. A decrease in phytochrome levels is expected to result in taller plants. A disruption in porphyrin metabolism by blocking MgChl is likely to result in an accumulation of PPIX to be available to ferrochelatase for producing heme, a precursor for the chromophore of phytochromes. An overproduction of phytochrome results in shorter Arabidopsis plants (Wagner *et al*. 1991). Thus, a simple flux increase towards the heme pathway would not be expected to increase plant height. Feedback regulation, due to PPXI accumulation, on the other hand, might restrict flux upstream in the porphyrin pathway and prevent the formation of bilin and perhaps heme. If this is the case, we expect that metabolite profiling experiments should demonstrate reduced accumulation of bilin, heme, PPXI, and upstream porphyrin intermediates in the *Oy1-N1989*/+ mutants. We expect plant elongation responses in *Oy1-N1989*/+ mutants to show reduced sensitivity to red and far red light. Still, we did not observe any such changes in the mesocotyls (Khangura *et al*. 2020b), although these do not accumulate high levels of chlorophyll and may not be as impacted by the *Oy1-N1989* mutant as in *elm1-ref*. In this study, we also found that maximizing elongation by disrupting bilin production with the *elm1* mutant did not increase plant height in *Oy1-N1989* and instead enhanced the loss of chlorophyll phenotype and dramatically decreased mutant plant height (**Figure 7**). Interpretation of this result is complicated by the decrease in chlorophyll levels, as a severe reduction of chlorophyll is expected to result in shorter mutant plants. One explanation for this observation may be upstream feedback regulation in porphyrin biosynthesis by bilin and phytochrome. Further work exploring porphyrin metabolism in these materials is necessary to test this.

The flowering time candidate gene analysis suggests another possible mechanism for the plant height effects of *Oy1-N1989*. The GA signal transduction transcriptional repressors, *d8* and *d9*, encode alleles that affected plant height in the mutant and mutant-to-wildtype height ratios (**Supplemental Table S12**). This suggests that GA signaling might be the mechanism responsible for the effect of *Oy1-N1989* on plant height. How chlorophyll reduction could result in a change in GA levels or sensitivity is unknown, but a test of this should be feasible. Similar to the experiments done with light and *elm1*, we expect that if GA is responsible for the increased height of *Oy1-N1989*/+, it should be sensitive to GA application, and the positive effects of *Oy1-N1989*/+ on plant height should be blocked in *D8/+;Oy1-N1989*/+ and *D9/+;O1-N1989*/+ double mutants.

### Representing allele effects in a molecularly interpretable manner improves candidate gene discovery for GWAS and QTL experiments

We and others have previously reported a QTL for plant height in B73 x Mo17 biparental populations co-localized with *zmmads69* and *zmsbp22* (Liu *et al*. 2015; Khangura *et al*. 2020b). In this study, we identified a SNP association linked to *zmmads69* and *zmsbp22* that impacted height and reproductive maturity traits in both mutant and wildtype siblings (**Figure 3a; Supplemental Table S5**). Increased expression of either of these two genes accelerated flowering and reduced plant height (Liang *et al*. 2019; Liu *et al*. 2020). Both are equally good candidates at this resolution permitted by RIL mapping. If mutants of only one had been assessed, that gene would be reported as a candidate gene “confirmed” by mutant studies. But the resolution provided by GWA and the additional data on gene expression allowed us to explore expression level polymorphism as a possible mechanism for this QTL. The direction of the natural variant’s allelic effect on the phenotype and proposed molecular mechanisms of the candidate gene must be considered to test this hypothesis. We previously eliminated a GA biosynthetic gene *dwarf plant3* as a possible cause of plant height variation reported by other labs using an inconsistency between an eQTL and plant height effect direction in the B73 x Mo17 population (Khangura *et al*. 2020b). We propose that allelic effects and molecular mechanisms be formally considered in evaluating candidate genes, when possible, to permit greater bridging between natural variant exploration and mechanistic studies. Such formal hypothesis construction and testing can help minimize the effects of the ascertainment bias that results from “confirming” candidates at genes for which we know the functions (Baxter 2020), a fallacy often compared to only searching for one’s lost keys under the lamppost where one can see at night.

In the current study, we compared our prior QTL analyses in B73 x Mo17 (Khangura *et al*. 2020b) to eQTL data from the same cross (Li *et al*. 2013, 2018). We also compared the eGWA data (Kremling *et al*. 2018) to the phenotypic trait GWA in the maize diversity panel. In the B73 x Mo17 biparental data, the expression of ZMMADS69 was increased, and ZMSBP22 was decreased by the allele associated with a reduction in plant height (**Supplemental Table S9**). This eliminates the eQTL at ZMSBP22 as a possible candidate for the plant height and flowering time QTL in B73 x Mo17 populations at this locus. We also detected a cis-eQTL in leaf tissue for ZMMADS69 (**Supplemental Figure S1; Supplemental Table S7**). Thus, the expression of ZMMADS69 was consonant with the phenotypic effect, suggesting that this locus could encode an eQTL affecting morphological trait variation.

We matched the allelic effects of SNPs at this locus on plant height, reproductive maturity, and expression to determine if they were under shared genetic regulation (**Figure 3; Supplemental Table S3**). Reproductive maturity and plant height were always affected in the same direction. Still, the accumulation of ZMMADS69 transcript was higher and lower, indicating recombination between the allele(s) affecting gene expression and the allele(s) affecting plant morphology and development. Thus, an eQTL is ruled out, and correlation due to linkage, not causation, is responsible for the relationships between expression and morphology (**Supplemental Figure S2; Supplemental Table S8**). This single-nucleotide resolution testing of allelic effects is a robust method for distinguishing possible molecular mechanisms. Further insight into the molecular mechanism by which the QTL at this position affects gene expression, height, and reproductive maturity will require allele-specific manipulation of plants using the natural alleles and protein-coding sequences.

The molecular nature of an allele will be the same regardless of the population in which it occurs or is measured. As such, it is possible to use molecular data to predict allelic effects on phenotype across populations. As long as the genetic basis for trait variation is the same and detectable, allele directions can be used to assess whether the same allele is being measured in two distinct populations. For example, the effect of the reference versus non-reference allele at a biallelic SNP is expected to remain the same regardless of the population. But using allele effects faces three basic technical limitations. First, natural variation may not be affected by simple biallelic mendelian phenomena. The complex linkage between multiple causative polymorphisms can create difficulties in GWAS. When variants of similar effects are linked in cis, they can form the basis of large-effect QTL in biparental populations where linkage is much more extensive (Flister *et al*. 2013; Khangura *et al*. 2019; Liu *et al*. 2020). The second limitation is one of allele frequency. In biparental populations, all alleles segregate ~at roughly 1:1, and we have maximum power to detect differences. In association mapping, rare alleles in the population will not be detected; indeed, we threshold SNPs below a frequency of 5% to avoid spurious associations and have no association results for these low-frequency positions. As a result, all alleles that are rare in an association population but present in either of the parents of a biparental population will contribute to the detection of QTL but be undetectable in an association experiment. The third limit is an artifact of the methods used for association mapping. This results from poor communication between computational biologists, statistical geneticists, and molecular biologists. The conventions of the most frequently used software tools for GWAS do not readily permit molecular hypothesis generation from these data. To achieve greater computational efficiency, SNP data are numericalized, and converted from nucleotides to numbers. The software tools used for this purpose can accomplish this goal differently. For example, in the GAPIT software version 3 (manual updated March 12, 2022; https://zzlab.net/GAPIT/), the default for this computational step results in allele effect directions oriented by the alphabetical order of the SNP alleles: “the sign of the allelic effect estimate (in the GAPIT output) is with respect to the nucleotide that is second in alphabetical order.” An alternative option, which is the default in software package TASSEL5 (Bradbury *et al*. 2007), permits selecting the most frequent allele in the population, the major allele, as the reference and the less frequent being coded as the alternative. This results in the sign of the allelic effect estimate being calculated with respect to the minor allele. That manual states, “In this scenario, a positive allelic effect estimate will indicate that the minor allele is favorable.” There is nothing inherently wrong with either of these options in a GWAS experiment, but when it comes to comparative analyses, overlooking how this step was carried out can create inconsistent allele effect directions among different tools for GWA and QTL mapping. The more problematic of the two is the major/minor approach. Suppose the major/minor allele approach is taken. In that case, variation in allele frequency affected by population sampling can result in an allele switching from being the major allele (e.g., allele frequency = 0.501) to being the minor allele (e.g., allele frequency = 0.499) and with this, a flip in the effect direction at such a SNP would be returned for two subsets of the same population by the same software package. As a result, downstream comparisons where the same QTL is compared across different approaches (GWA and QTL mapping; between two GWA experiments using different tools; even between two GWA experiments using the same tool and different populations or subsamples from the same starting population!) can return opposing allele effect signs in the output. This prevents biological interpretation that uses experimental comparisons at nucleotide resolution. Using an established reference genome and alternative alleles to orient the SNP coding ties the effect direction from the GWA analysis to the molecular nature of the mutation. This makes mechanistic follow-up and communication between population genetics and molecular genetics possible.

Because we wanted to compare our GWAS results to our B73 x Mo17 QTL results, we numericalized our SNP data such that all B73-reference alleles were coded as the reference and effect directions were calculated relative to the effect of the non-B73 allele. This was accomplished by replacing all the B73 reference alleles in our genotype VCF files with “A” and all the non-reference alleles with “T” to fool the alphabetical numericalization implemented in GAPIT to generate a numericalized SNP file that allows us to make molecular predictions about allelic effects relative to the reference genome by looking at the allele sequence. This conversion provides the SNP alleles to perform analysis such that the sign of β (*beta*, used to denote allele effect) can be compared to our previous QTL mapping results that compared the allelic effects of B73 and Mo17. It is worth noting that co-localization of a QTL from different populations (e.g., biparental and association mapping panels) does not guarantee a shared molecular basis. For example, out of the five QTL-GWA co-localizations in our work, only one, *oof1* (Khangura et al., 2020a), failed to show a consonant effect direction. Thus the co-localization of QTL and GWAS was spurious. The other four have consonant effects of the alternate allele in both QTL and GWAS experiments, and three have excellent candidate genes (**Supplemental Table S15**). We propose that as more GWA experiments are conducted and attempts are made to derive molecular identities for the natural alleles discovered, calculation of allele effects relative to a reference genotype (e.g., B73 for maize, Columbia-0 for Arabidopsis, etc.), or a clear explanation on how effect directions were calculated will help bridge the quantitative-molecular divide and accelerate discovery. Our numericalized SNP file and a dummy SNP file that reorients the SNPs from maize HapMap3 (Bukowski *et al*. 2018) are available with this manuscript. We encourage researchers working on other organisms to use a similar approach when comparing alleles across experiments.

## Supplemental Figures

**Supplemental Figure S1.**
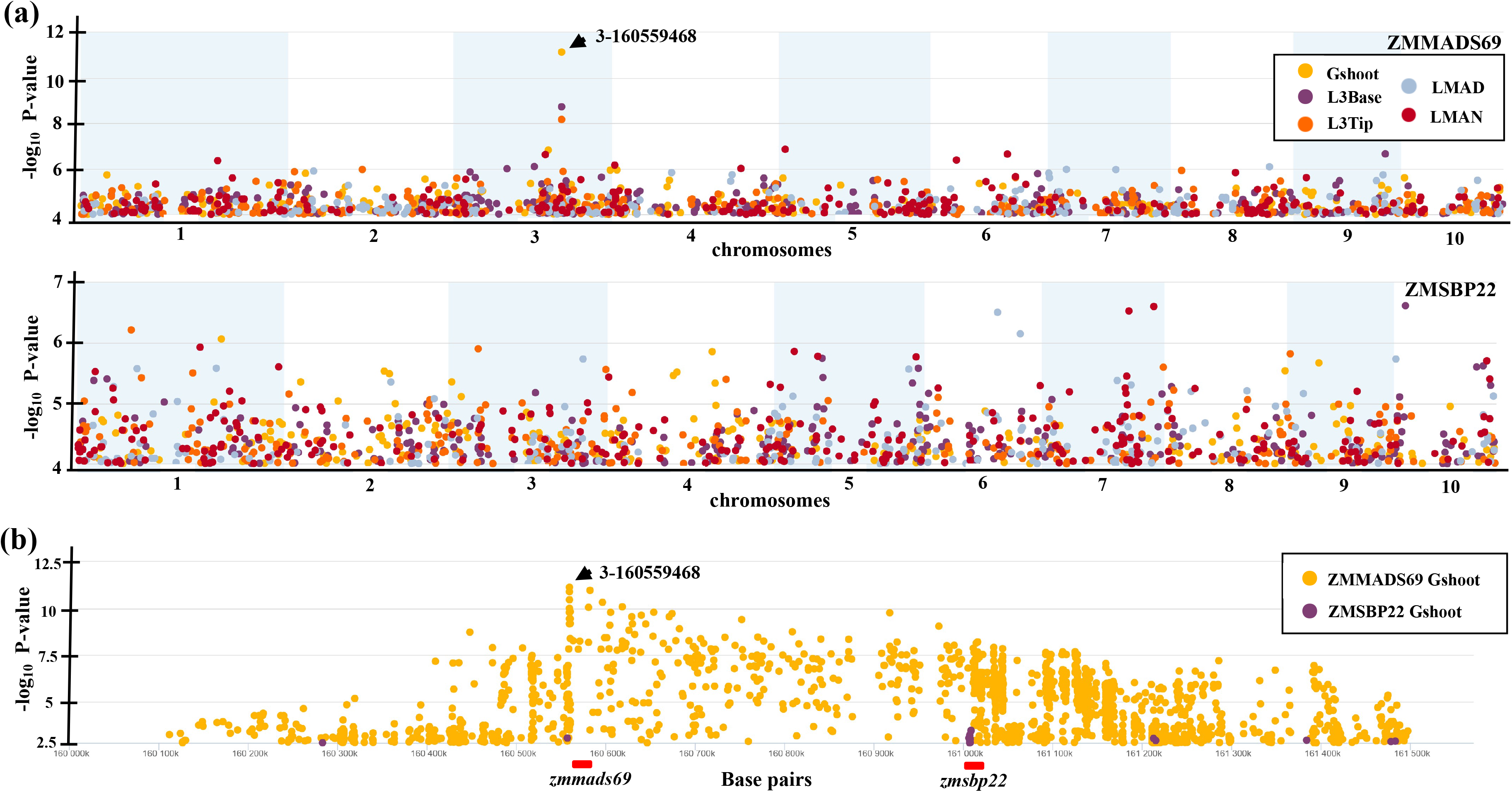
Manhattan plot displaying the association mapping of accumulated transcripts in five green tissues for (a) ZMMADS69 and ZMSBP22. Only SNPs with P-value below 10^−4^ in a 250 kb window are plotted. (b) A 1.4 Mb window showing cis-acting SNP association with ZMMADS69 and ZMSBP22 in germinating maize shoots. The red bar underneath the X-axis shows the relative position of *zmmads69* and *zmsbp22* gene models.

**Supplemental Figure S2.**
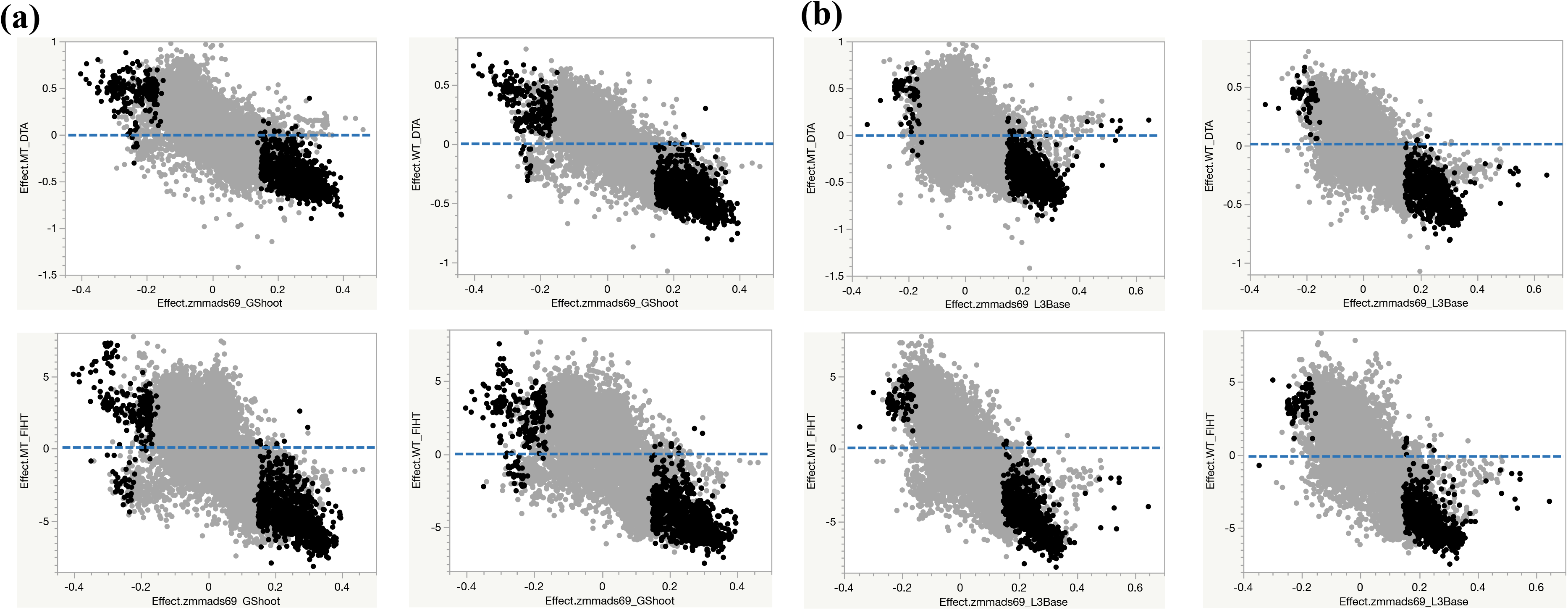
Bivariate plot of allele effects of *zmmads69* transcript accumulation in different tissue in the inbreds in the maize association panel (X-axis) and the F1 hybrids of the Oy1-N1989/+:B73 with the association panel (Y-axis) for mutant days to anthesis, mutant flag leaf height, wildtype days to anthesis, and wildtype flag leaf height. Black dots represent SNPs with P-value < 0.001 for association analysis using *zmmads69* transcripts in (a) germinating shoots (Gshoot), and (b) base of third leaf (L3Base). Blue dotted line is drawn to highlight SNPs with low expression and high trait effect by the alternate allele and vice-versa.

**Supplemental Figure S3**. Bivariate plots of effects of alternate allele on ZMMADS69 transcript accumulation in the tip of third leaf in the inbreds in the maize association panel (X-axis) and the F1 hybrids of the *Oy1-N1989*/+:B73 with the association panel (Y-axis) for mutant days to anthesis, mutant flag leaf height, wildtype days to anthesis, and wildtype flag leaf height.

## Supplemental Tables

**Supplemental Table S1**. The BLUP values of all the traits in the F1 hybrids between the maize diversity panel and the *Oy1-N1989*/+:B73 tester.

**Supplemental Table S2**. Pearson pairwise correlations between various traits measured in the F1 hybrids between the maize association panel and *Oy1-N1989*/+: B73 tester.

**Supplemental Table S3**. Genome-wide SNP associations that were significant at Bonferroni-corrected genome-wide threshold in the F1 hybrids between the maize diversity panel and the *Oy1-N1989*/+:B73 tester.

**Supplemental Table S4**. Effect direction of the top vey1 SNPs for all wildtype and *Oy1-N1989*/+ mutant traits in the F1 hybrids relative to the *vey1* effect.

**Supplemental Table S5**. Genome-wide statistical significance and effect of all traits analyzed in this study at SNP positions with any Bonferroni-corrected association for any given trait.

**Supplemental Table S6**. Result of the eQTL mapping for ZMMADS69 and ZMSBP22 transcript accumulation in the maize shoot apex of IBM RILs.

**Supplemental Table S7**. Test statistics for 14294 SNPs in the 1.4 million base pair region around *zmmads69* and *zmsbp22* for its transcript accumulation in the maize association panel and traits from the F1 population between the maize association panel and *Oy1-N1989*/+:B73 tester.

**Supplemental Table S8**: List of SNP associations that passed the genome-wide Bonferroni threshold after using three top SNPs at *oy1* as covariates in the model.

**Supplemental Table S9**. Candidate genes from Arabidopsis and their orthologs in maize in the porphyrin, heme, siroheme, and chlorophyll biosynthesis pathway used for pathway level analyses.

**Supplemental Table S10**. SNPs that yielded significant associations at p<10^−4^ with linked enzymes in porphyrin, heme, siroheme, and chlorophyll biosynthesis pathway.

**Supplemental Table S11**. List of *bona fide* regulators of flowering time in maize obtained from literature search.

**Supplemental Table S12**. SNPs that yielded significant associations at p<10^−4^ with linked genes that are *bona fide* regulators of flowering time in maize.

**Supplemental Table S13**. QTLs and candidate genes for each locus detected in the IBM x *Oy1-N1989*/+:B73 and Syn10 x *Oy1-N1989*/+:B73 F1 mapping populations described previously.

**Supplemental Table S14**. QTLs from previous studies and GWA SNP associations from this study that co-localized and had a consonant allelic effect on the trait identifies candidate genes.

**Supplemental Table S15**. Morphometric analysis of single and double mutants between *Oy1-N1989*/+ and *elm1-ref* in the B73 inbred background.

## Acknowledgements

The authors would like to acknowledge the assistance of the staff and leadership at the Agronomy Center for Research and Education and the Horticulture Greenhouse at Purdue University. We are grateful to Dilkes lab members for comments and discussions, particularly Dr. Amanpreet Kaur for her critical reading of drafts. We would particularly like to thank everyone who puts extra effort into scientific software documentation that allows research such as this to determine the meaning of all values output from open-source resources. You are the real heroes. This work was supported by United States Department of Agriculture National Institute of Food and Agriculture postdoctoral grant 2022-67012-36601 to RSK and United States Department of Energy Office of Science (BER) Grant DE-SC0023305 to BPD.

## Notes

**Conflict of Interest** Authors declare no conflict of interest.

